# Abnormal nuclear organization impairs adipogenic lineage commitment and terminal maturation in Hutchinson-Gilford progeria syndrome (HGPS)

**DOI:** 10.64898/2026.10.19.760205

**Authors:** Nishita Gogia, Alvaro Rodriguez Gonzalez, Lily Eisenklam, Rebeca San Martin, Rachel Patton McCord

## Abstract

Hutchinson-Gilford progeria syndrome (HGPS) is a fatal premature aging disorder driven by progerin, a truncated Lamin A protein that destabilizes nuclear architecture and disproportionately affects mesenchymal-lineage tissues. One notable mesenchymal lineage defect in Progeria patients is early and severe subcutaneous lipodystrophy. We hypothesized that disorganized chromatin architecture underlies the failure of mesenchymal stem cell (MSC) lineage commitment and adipogenesis in HGPS. To test this while preserving donor age, epigenetic history, and tissue context, we induced adipogenesis in age-matched healthy and HGPS patient-derived dermal fibroblasts and profiled cells using imaging, bulk RNA-seq, and Hi-C across the differentiation time course. We found that HGPS fibroblasts commit to the adipogenic lineage at rates comparable to healthy controls but diverge downstream: induced cells display aberrant lipid-droplet morphology, mount a disorganized and delayed transcriptional response, and retain priming for alternative mesenchymal lineages (osteochondrogenic, vascular) alongside stalled adipogenic maturation. Hi-C revealed that HGPS fibroblasts exhibit some baseline spatial compartment differences from healthy cells, but more notably show differences in how they reorganize compartments during adipogenic induction. Compartment-transcription coupling was strong during adipogenesis for lipid-metabolic genes in healthy cells but was more dysregulated in HGPS. These results indicate that HGPS disrupts lineage consolidation rather than initiation, associated with disease-specific failures in chromatin reorganization during differentiation.

## Introduction

Hutchinson-Gilford progeria syndrome (HGPS) is a rare disease characterized by a severe premature aging phenotype [1, 2]. It is caused by a single-point mutation in the Lamin A gene (LMNA, c. 1824C>T). This mutation causes striking features of rapid, premature aging in children, with an average life expectancy of about 14.5 years [3, 4]. The mutant Lamin A activates a cryptic splice site within exon 11 [5, 6]. The resulting transcript encodes a mutant Lamin A protein called progerin; this variant lacks the internal ZMPSTE24 cleavage site required for Lamin A maturation; progerin therefore retains the C-terminal farnesyl anchor and permanently tethers to the inner nuclear membrane [7]. Progerin acts in a dominant-negative manner, leading to nuclei becoming lobulated/blebbed, mechanically stiff, and destabilizing the contacts between the lamina and the underlying chromatin, ultimately leading to deterioration of genome organization in the affected cells [8, 9]. At the tissue level, patients present with short stature and a slow growth rate, marked lipodystrophy with loss of subcutaneous fat, osteoporosis, and progressive arterial stiffening [6, 10, 11]. Notably, nearly all the affected tissues are derived from the mesoderm, more specifically the mesenchymal lineage, pointing toward an upstream dysregulation of the mesenchymal stem cell (MSC) lineage commitment rather than a defect confined to terminal cell and tissue types [12–16].

Whether the architectural defect and the mesenchymal tissue phenotype are causally linked to one another remains untested; therefore, we hypothesize that disorganized chromatin is the mechanism by which progerin disrupts mesenchymal lineage commitment, making the fat, bone, cartilage, and vasculature phenotypes downstream consequences of a single architectural defect rather than independent tissue failures.

Of particular interest to us is the loss of subcutaneous fat, which is both drastic and early in these patients: lipodystrophy is detectable within the first year of life [17, 18]. Weight gain slows from around two years of age [19] and thereafter affected children fail to accrue body fat despite adequate caloric intake [10]. Laminopathies involving the LMNA gene have been broadly associated with defects in lipid metabolism and adipocyte differentiation [20], and in HGPS, the accumulation of progerin has been tied directly to impaired adipogenesis [21]. In vitro studies using iPSCs derived from patient-derived fibroblasts have shown that progerin represses activation of the core master regulators like PPARγ2 and C/EBPα, arresting adipocytes before terminal maturation [22]. Premature cellular senescence appears to compound this block: progerin-expressing senescent cells upregulate inflammatory cytokines that limit the proliferative expansion and differentiation potential of adipose progenitor cells [23].

What remains unclear is how these transcriptional deficits reported in the literature are encoded at the level of genome architecture. Addressing this directly in patients is not feasible, as the bone marrow biopsy is untenable given the juvenile mortality of the patients. Existing in-vitro cell culture models carry a common limitation: iPSCs [13, 15, 22, 24] and Skin-derived Precursor cells (SKPs) [23] based systems begin from a multipotent or reprogrammed state, stripped of local context, senescent neighbors, chronic inflammatory signaling, and other niche-derived cues that shape the adipogenic decision in the patients.

We therefore use age-matched healthy and HGPS patient-derived dermal fibroblasts. These retain the donor’s age and epigenetic landscape, context that reprogrammed cells lose, while harboring an MSC-like population that permits the lineage commitment to be studied within a heterogeneous culture [25], which we refer to as an MSC-like subpopulation in the manuscript. This model system is further motivated by a previous finding that HGPS fibroblasts carry vestigial transcription signatures indicative of aberrant mesenchymal commitment [26], confirming that the lineage deficit is present in this system.

To determine whether this signature reflects an underlying defect in the genome organization, we combined high-throughput chromatin conformation capture (Hi-C) with transcriptomic profiling and cellular phenotype imaging across adipogenic induction, resolving how high-order chromatin folding is remodeled or fails to remodel as HGPS cells attempt commitment towards the adipogenic lineage.

Our results indicate that HGPS cells commit to the adipogenic lineage in numbers comparable to controls but diverge downstream of that decision. Induced progeria patient cells yield adipocytes of aberrant morphology, mount an exaggerated transcriptional response in both breadth and magnitude to adipogenic stimulus, and retain 3D genome structural priming of loci belonging to alternative mesenchymal lineages while the adipogenic program is stalled in its initial state. We propose that this failure to restrict lineage development potential, rather than a failure to initiate commitment, underlies the loss of subcutaneous fat that characterizes these patients.

## Results

### Adipogenic competence and remodeling are separable in MSCs and fibroblasts, and progeria affects only the latter

In the literature, it is reported that MSCs and fibroblasts share a surface marker profile, making the two cell types difficult to distinguish [27, 28]. We therefore asked whether the MSC subpopulation within the fibroblast culture remodels towards an adipogenic fate in the same manner as bone marrow MSCs. To address this, we compared adipogenic induction in bone marrow MSCs and 8-year-old healthy fibroblasts (GM08398, called “08398” hereafter) at day 7 (D7) (Supplementary Figure 1A) and day 21 (D21) (Figure 1B).

**Figure 1:**
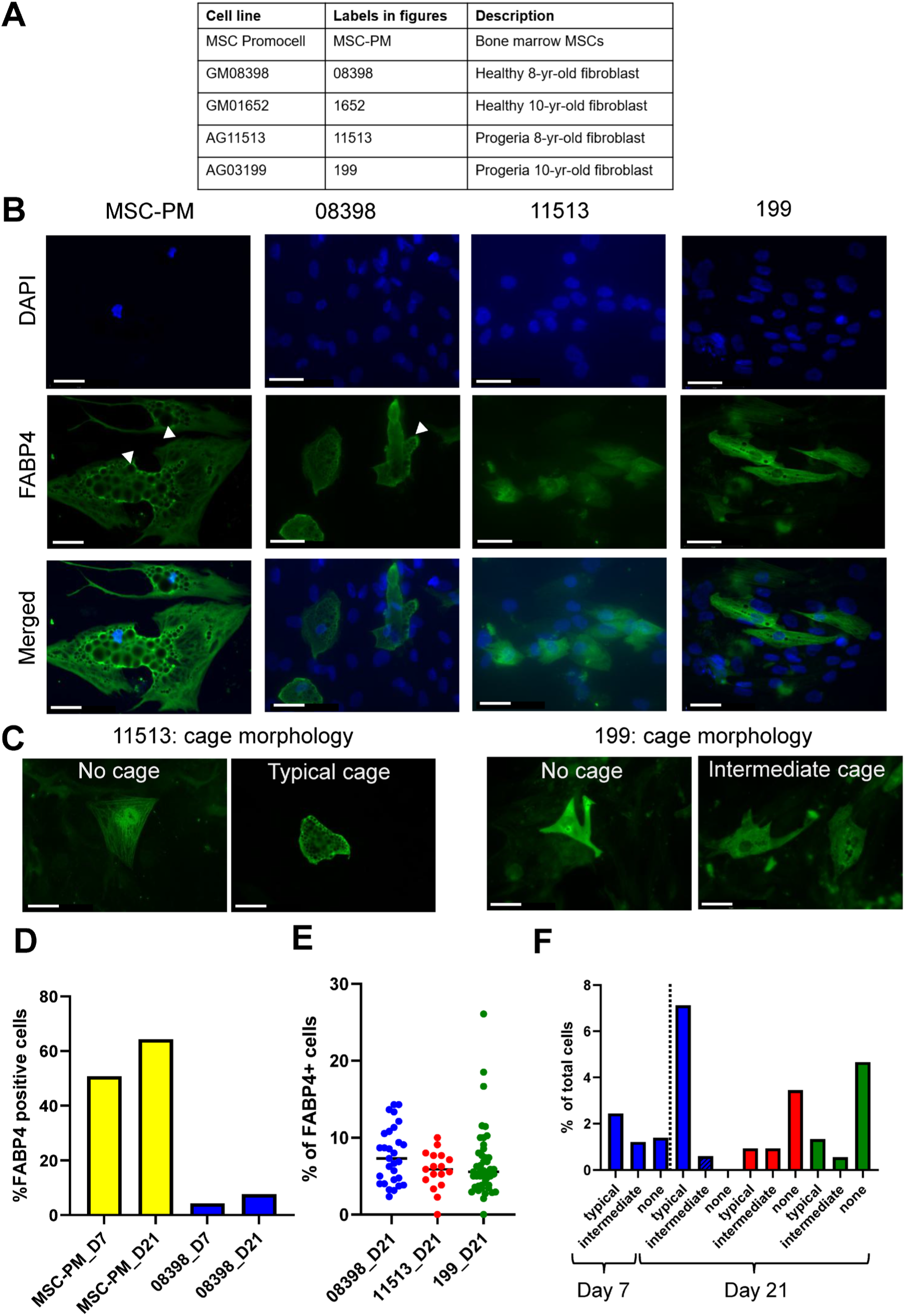
Progeria fibroblasts show normal adipogenic commitment but heterogeneous, defective cage remodeling. A) Cell lines used in this study with the labels used in the figure and a description of each. (B) Representative immunofluorescence images of bone marrow MSCs (MSC-PM), healthy 8-year-old fibroblasts (08398), and progeria fibroblasts (11513 and 199) at day 21 of adipogenic induction, stained for FABP4 (green) and nuclei (DAPI, blue), Scale bars: 50 µm. (C) Representative images of cage morphologies in FABP4+ progeria cells at day 21, showing cells without cages and cells with typical or intermediate cage-like structures in 11513 (left) and 199 (right). Scale bars: 50 µm. (D) Percentage of FABP4+ cells in MSC-PM and 08398 fibroblasts at day 7 (D7) and day 21(D21) of adipogenic induction. Each bar represents cumulative number of FABP4+ cells per field captured. Number of nuclei in each sample (n): MSC-PM D7= 65, MSC-PM D21= 42, 08398 D7 = 984, 08398 D21 = 828. (E) Percentage of FABP4+ cells in healthy (08398) and progeria (11513 and 199) fibroblasts at D21. Each dot represents one imaging field/well; horizontal lines indicate the mean ± SEM. Number of nuclei in each sample (n): 08398: n= 828, 11513: n= 319, 199: n= 1264. Differences between groups were not significant (Kruskal-Wallis test, H=3.28, p=0.19). (F) Quantification of cage morphology (typical, intermediate, none) in FABP4+ cells, expressed as a percentage of total differentiated cells. Healthy fibroblasts (blue) are shown at D7 and D21; progeria lines 11513 (red) and 199 (green) are shown at D21.

Induction efficiency and commitment to the adipogenic state, quantified as the fraction of FABP4+ cells, differed markedly between the two cell lines: MSCs committed 64% of the population to adipocytes, compared to only 7% for the 08398 fibroblasts (Figure 1D). The difference was not limited to competence but also differed in morphology. MSC-derived adipocytes were larger overall and displayed more prominent “cage-like” structures characteristic of typical adipogenesis, as reported in the literature before [29, 30] whereas the equivalent structures in the fibroblast MSC subpopulation-derived adipocytes were smaller (Figure 1B, marked by white arrows), suggesting a difference in remodeling during adipogenic induction between bone marrow MSCs and the MSC subpopulation within fibroblast culture. Both lines showed stronger induction and more advanced commitment to D21 than on D7 (Figure 1D), and we therefore restricted the remainder of our analysis to D21.

Across all cell lines, progeria and their healthy age-matched controls, the percentage of FABP4+ cells did not differ significantly (Figure 1E), indicating comparable adipogenic potential by this measure. However, morphological analysis revealed heterogeneity in the cage-like structures among the progeria cells that were absent from the controls. These cage-like structures likely arise when vimentin filaments enclose the lipid droplets formed during adipogenesis [30]. To establish whether lipid droplet filling and cage assembly are coupled events, we manually inspected all images for their co-occurrence. The two events were not linked in all instances; cage formation and lipid droplet formation could proceed independently of one another (Supplementary Figure 1B).

We therefore categorized the cage morphology of FABP4+ healthy and progeria cells into three classes: cells with typical cage-like structures, cells with fewer and smaller cages (“intermediate”); and cells in which cage morphology was absent altogether (Figure 1B and 1C). In healthy controls, cells increasingly showed typical cage-like structures from the D7 to D21 timepoints, accompanied by a decline in intermediate structures consistent with intermediate structures progressively resolving into mature cages (Figure 1F). This suggests that healthy donor cells can resolve their cage architecture over time as they reach a fully committed adipocyte state. In both progeria lines, by contrast, the majority of FABP4+ cells had no cages, rather than typical morphology, even at 21 days of differentiation.

Together, these results indicate that the adipogenic deficit in progeria patient-derived fibroblasts is not a matter of how many cells respond to differentiation cues, but of how they remodel once committed. FABP4 induction rates are like age-matched controls, but cage morphology diverges sharply, a dissociation that refines the standard interpretation of the poor droplet formation (Supplementary Figure 1B) and subcutaneous fat loss documented in HGPS.

Two aspects of this phenotype, to our knowledge, have not been described previously. First, cage assembly and droplet filling are separable events, so the cage phenotype cannot be dismissed as a downstream effect of the known droplet deficit. Second, adipogenic induction in progeria cultures results in a mixture of typical, reduced, and absent cage morphologies rather than a uniform defect. The progeria phenotype is therefore one of heterogeneous remodeling rather than uniform failure of the adipogenic program.

### MSCs and fibroblasts are transcriptionally distinct, and progeria fibroblasts show pathway-selective loss of stemness

Building on the FABP4 quantification, which showed bone marrow MSC, healthy fibroblast, and progeria fibroblast cultures differ in their adipogenic response, we next asked how stem-cell marker expression differs between these cell types and how it shapes their adipogenic trajectories. To compare the transcriptional response to adipogenic induction in MSCs vs the healthy patient-derived fibroblast culture, we used published bulk RNA-seq data from umbilical cord MSCs [31], sampled at Day 0 (D0) and day 15 (D15) of adipogenic differentiation. For the patient-derived fibroblasts, we performed bulk RNA-seq from four donor cell lines: two healthy donors, one 8-year-old (08398) and one 10-year-old (1652), and two age-matched HGPS patients carrying the canonical LMNA c.1824 C>T mutation, one 8-year-old (11513) and one 10-year-old (199). For each donor, cells were sampled at three timepoints: day 0 (D0, uninduced control), day 7 (D7), and day 21 (D21) under both control and adipogenic induction conditions. This design enabled us to compare age-matched healthy and progeria donors at matched donor ages and differentiation stages, and to benchmark the fibroblast adipogenic trajectory against the MSC reference.

To characterize how MSCs vs fibroblast cultures remodel their transcriptional landscape during adipogenesis and how this remodeling is disrupted in HGPS patient-derived fibroblast cultures, we assembled a curated stem-cell marker list from publicly available databases (see methods: MSC vs fibroblast transcriptional profiling) and overlapped it with our RNA-seq dataset. Three distinct expression patterns emerge: (i) a set of seventeen markers were expressed in MSCs but silent across all fibroblast cell lines. Most were self-renewal/lineage signaling genes (e.g., CTNNB1, AGO2, GPC3); only one pluripotent marker, STAT3, followed this pattern, suggesting a canonical MSC-restricted signature. (ii) Eight mesenchymal and neural-progenitor markers, including MME(CD10), TGFBR2, FAT4, and SIX4, were expressed in MSCs and healthy fibroblasts but lost in both HGPS donors. (iii) Seventeen markers, including TMEM64, BMP8A, HNRNPU, HOXA11, SMAD4, and KANK1, were maintained specifically in the healthy fibroblasts and lost in HGPS, defining a fibroblast-specific stemness signature. (iv) HGF and ACVR2 were induced during adipogenesis in healthy fibroblasts but not in HGPS (Supplementary Figure 1C).

These findings corroborate our imaging results in that MSCs and fibroblasts wire their transcriptional landscape through distinct signaling pathways during adipogenic differentiation. The MSC subpopulation of HGPS fibroblast cultures selectively lose mesenchymal identity and fibroblast-specific programs rather than stemness globally. To understand how disease alters this fibroblast-specific adipogenesis, we compared healthy and HGPS donors’ responses to adipogenic induction over time.

### A temporal lag in the adipogenic window in HGPS: disrupted age-dependent timing and failure to consolidate an expanded transcriptional response

Among the healthy cells, 8-year-old donor 08398 exhibited more DEGs (D7 or D21 unique genes; see methods: RNA-seq analysis) than 10-year-old donor 1652 at early commitment, a pattern that reversed at terminal differentiation (Figure 2A; healthy donors). This suggests an age-stratified temporal shift, in which the younger donor initiates adipogenic commitment earlier than the older donor. To determine whether the 08398 D7 head-start genes were also activated in 1652 with delayed kinetics, we assessed how many of these D7 genes were upregulated in 1652 at D21. Of the 246 genes upregulated specifically in 08398 at D7, 24% (58/246) reached significance in 1652 by D21; all were concordant in direction, suggesting these genes were delayed in their response in the older donor (Supplementary Figure 1D).

**Figure 2:**
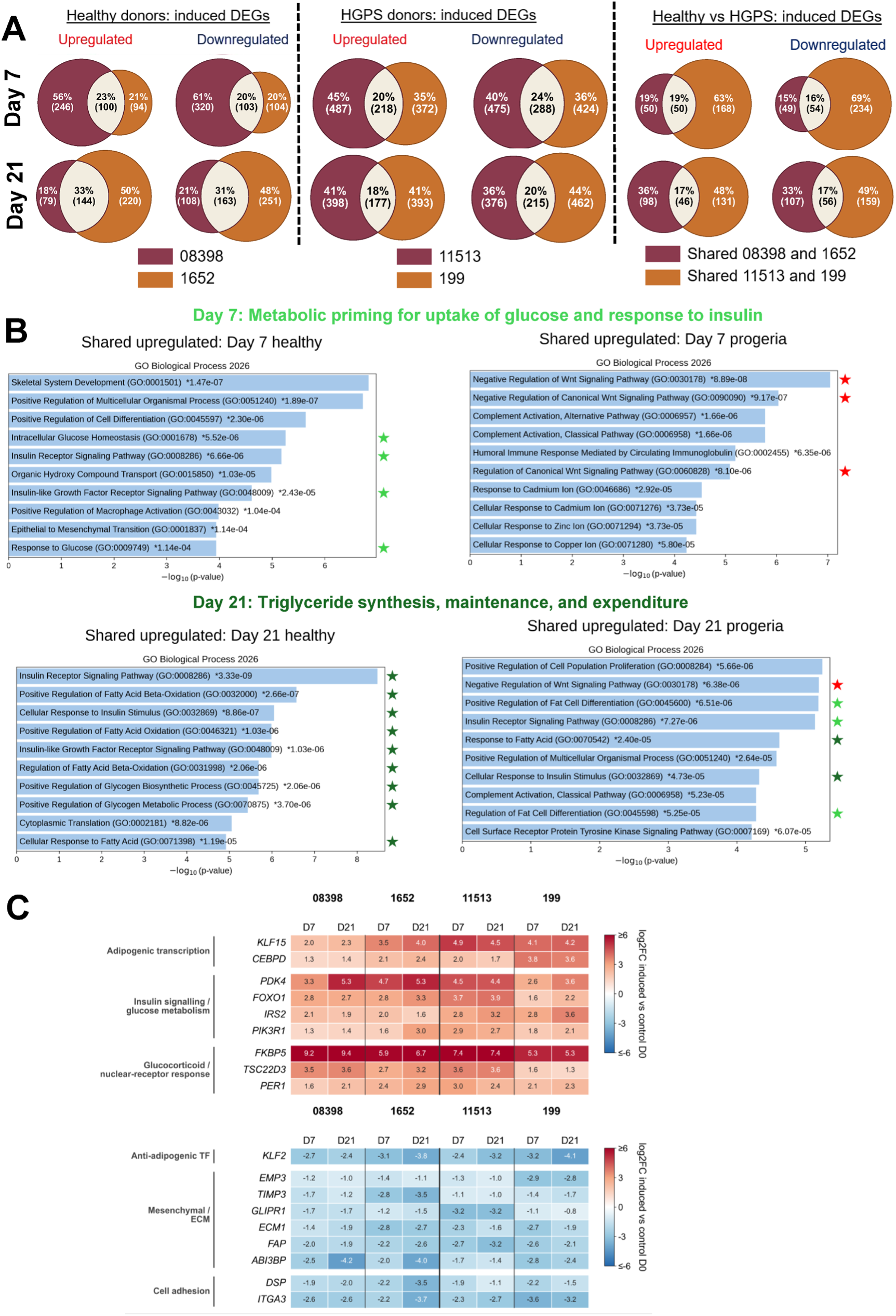
Healthy and progeria fibroblasts activate shared adipogenic program, though with distinct enriched pathways. (A) Venn diagrams of differentially expressed genes (DEGs) induced by adipogenic differentiation relative to undifferentiated control (D0), at day 7 (top) and day 21 (bottom). The left panels compare the two healthy donors (08398 and 1652). The middle panels compare the two HGPS donor lines (11513 and 199); the right panels compare the DEGs shared by both healthy lines with DEGs shared by both HGPS lines. Upregulated and downregulated DEGs are shown separately. Values are the percentage and number of genes in each donor. DEGs were defined as adjusted p < 0.05, as determined by DESeq2. (B) Gene Ontology (GO) Biological Process enrichment of the upregulated DEGs shared by both healthy lines (left) or both HGPS lines (right) at D7 (top) and D21 (bottom). The top 10 terms ranked by -log10(p-values), and the p-values for each term are shown next to its bar. Star terms related to early adipogenic signals involving glucose and insulin, late adipogenic signals like fatty acid oxidation and lipid metabolism with dark green, and inhibition of adipogenesis like Wnt signaling with red stars. (C) Heatmaps of selected induced genes in each cell line at D7 and D21, shown as shrunken log2fold change versus D0 controls. The top heatmap shows upregulated genes grouped by function: adipogenic transcription, insulin signaling and glucose metabolism, and glucocorticoid receptor response. The bottom heatmap shows downregulated genes: an anti-adipogenic transcription factor, mesenchymal/ECM genes, and cell adhesion genes. Each cell shows its log2FC values, and the color scale is capped at ±6 (biological replicates, n=2).

This age-dependent temporal shift was not recapitulated in the age-matched HGPS samples: both 11513 (8-year-old) and 199 (10-year-old) maintained equal DEG counts at both timepoints (Figure 2A; HGPS donors). Progeria samples consistently showed approximately 1.5-2.9-fold more DEGs than their age-matched healthy controls at both timepoints, in both directions (Figure 2A; Healthy vs HGPS and Supplementary Figure 3A), indicating more extensive, bidirectional transcriptomic remodeling during adipogenic differentiation in HGPS donors. Despite these differences, all four donors shared a core induction response: 60 genes changed in the same direction at both D7 and D21 in every donor (28 upregulated and 32 downregulated). Genes induced in all donors were enriched for adipogenic and insulin signaling GO terms, including white fat cell differentiation and insulin-receptor signaling (Supplementary Figure 2D). These terms were supported by the adipogenic factors CEBPD and KLF15 and the insulin-pathway genes IRS2, PIK3R1, FOXO1, and PDK4, as suggested by the GO terms. Glucocorticoid-responsive genes were also uniformly induced, consistent with adipogenesis programming (Figure 2C). Conversely, genes repressed in all donors were enriched for a mesenchymal/ECM program, and the anti-adipogenic factor KLF2, a repressor of PPARγ, was downregulated in every donor. This indicates that adipogenic induction suppresses fibroblast ECM programs and releases an inhibitor of adipogenesis even in HGPS cells (Figure 2C), which is also supported by the GO terms (Supplementary Figure 2E).

To determine which pathways are engaged at what stage in progeria cells, we compared genes shared between the two healthy donors with those shared between HGPS donors. About half of the genes changed in healthy donors at D7 are also changed in both HGPS patients at this time point, and 32-34% of DEGs are shared by D21 (Supplementary Figure 2A). When we compared the functional categories of these gene sets, we noted a temporal lag in HGPS adipogenesis pathways: In healthy donors, GO enrichment for the upregulated genes shifts across the induction window, moving from metabolic priming, glucose uptake, and insulin response at D7 towards fatty acid oxidation and glycogen metabolism by D21 (Figure 2B). In contrast, the HGPS shared set of upregulated genes on D7 includes no glucose, insulin, or fatty acid terms, but instead includes inflammatory and Wnt signaling pathways. By D21, the HGPS gene set functions resemble a mixture of the healthy D7 and D21 profiles, indicating that progeria cells reach the early stage of the adipogenic cascade late and abnormally. Notably, very few genes change in opposing directions in healthy vs. HGPS cells (Supplementary Figure 2A), suggesting that some genes fail to change in the right direction rather than misregulating in the wrong direction. This loss reflects erosion rather than redirection: across all donors, DEGs retain their direction of regulation.

Together, these results indicate that HGPS cells do not fail to initiate adipogenesis; rather, despite broader transcriptional remodeling, the adipogenic program is engaged but with delayed timing, and fails to consolidate.

To move beyond broad functional enrichments in full gene sets to more specific examination of relevant pathways and genes, we curated functional panels spanning compromised pathways reported in the HGPS literature, like Wnt signaling [32–34], cell cycle [34–36], oxidative stress, skeletal and vascular programs, and adipocyte markers [10], comparing each HGPS donor against its age-matched healthy control (see methods; Pathway-level log2FC plot). We take the panels in turn, noting for each where the HGPS donors behave alike and where they diverge.

### Mis-coordinated adipogenic identity in HGPS: dysregulated PPARγ target effectors, mis-timed Wnt inhibition and mesenchymal drift

#### Adipocyte markers and PPARγ targets

PPARγ-target effectors play a central role in the regulation of lipid and glucose homeostasis during adipogenesis [37, 38]. In 11513, these effectors were selectively dysregulated (Figure 3A; adipocyte markers): genes like LPL, LEP, and CIDEC were induced from D7, while 08398 induced none of these three, and FABP4 rose further by D21, where it attenuated in 08398. In 199, the same panel was attenuated, with FABP4 and LPL falling below 1652 at D21; CIDEC is never induced, and LEP appears earlier than in 1652, at D7 rather than D21. This parallels the imaging data, where the FABP4+ cells accumulated in 11513 but with abnormal morphology in comparison to 08398, suggesting that HGPS vs. healthy donors consolidate the adipogenic program differently by D21.

**Figure 3:**
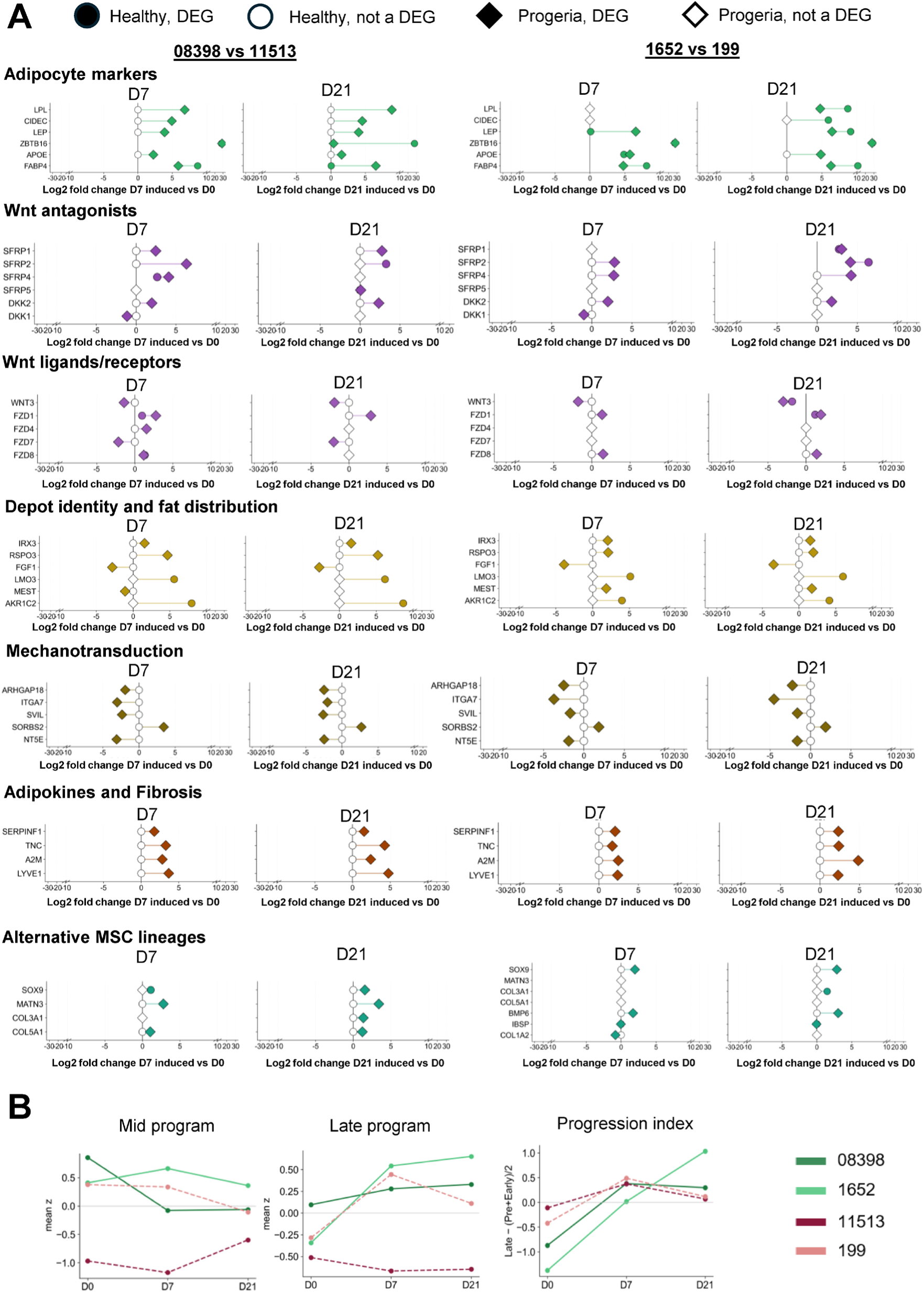
Progeria fibroblasts show altered induction of adipogenic, Wnt, and lineage-related genes during adipogenic differentiation. (A) Dumbbell plots comparing the log2fold change of selected genes at D7 and D21 of adipogenic induction, relative to undifferentiated controls (D0), in each healthy line and its age-matched progeria line: 08398 vs 11513 (8-year-old donors, left) and 1652 and 199 (10-year-old donors, right). Circles represent healthy lines and diamonds represent progeria lines. Filled symbols mark genes that are differentially expressed in that line. Open symbols mark genes that are not differentially expressed and are plotted at log2FC=0. Values are apeglm-shrunken log2fold changes;DEGs were defined as adjusted p <0.05, so a few significant genes have shrunken log2FC close to 0. A line connects the healthy and progeria values for each gene. Genes are grouped by function: adipocyte markers, Wnt antagonists, Wnt ligands and receptors, depot identity and fat distribution, mechanotransduction, adipokines and fibrosis, and alternative MSC lineages. The x-axis is compressed beyond |log2FC|= 10. (B) Gene modules scored across adipogenic differentiation (D0, D7, D21) for healthy (08398 and 1652, green solid lines) and progeria (11513 and 199, red dashed lines) fibroblasts. Expression values (log2CPM) were averaged across replicates (n=2) and z-scored per gene across all donors and time points. Mid and late program scores are the mean of z-scores of replicates (n=2) of genes in each module. The progression index is calculated as late – (Pre + Early)/2. Module gene lists are provided in Methods: Per-stage RNA-seq and Hi-C chromatin trajectory analysis.

#### Wnt antagonists, ligands and receptors

Another important axis for adipogenesis to proceed in an orchestrated manner is the timing of the Wnt-inhibitory checkpoint [39]. The secreted Wnt antagonists SFRP2, SFRP4 and DKK2 are induced at D7 in both HGPS donors and SFRP1 at D7 in 11513 and D21 in 199 (Figure 3A; Wnt antagonists), whereas their age-matched controls induce these genes later or not at all: at D7, only 08398 induced SFRP4; by D21 08398 induced SFRP2 and 1652 induced SFRP1 and SFRP2; and neither healthy donor induced DKK2. This later induction in healthy cells may reflect a feedback mechanism that restrains adipogenesis at that stage. At D21, 199 sustains all four antagonists, whereas 11513 only retains SFRP1 and DKK2,

Wnt ligand and receptor changes followed the same early timing. Both HGPS donors repressed WNT3 and induced FZD1 at D7, whereas 1652 made these changes only at D21; 08398 also induced FZD1 at D7. In addition, 11513 induced FZD4 and repressed FZD7, and 199 induced FZD8 (Figure 3A; Wnt ligands/receptors). Together, this data suggest that HGPS cells can engage Wnt inhibition but do so earlier than healthy cells.

#### Depot identity and fat distribution

Both healthy donors induce LMO3, a LIM-domain transcriptional cofactor linked to glucocorticoid receptor signaling in adipose tissue, and AKR1C2, important for sterol/lipid metabolism at D7 and D21, and neither HGPS donor does; conversely, both HGPS donors induce IRX3 and RSPO3 and repress FGF1 at both time points, and neither healthy donor does. IRX3 is the FTO effector that shifts adipocytes away from thermogenesis toward lipid storage, and FGF1 is a PPARγ target required for adipose remodeling. The HGPS lines therefore acquire storage drive while losing the capacity to remodel the depot that would hold it, a combination predicted to yield poorly expandable, fibrotic fat rather than more fat and one that sits with the fibro-adipokine signature described below (Figure 3A).

#### Mechanotransduction and MSC identity

Both HGPS donors repress the RhoA inactivator ARHGAP18, the laminin receptor ITGA7, and the actin membrane linker SVIL, and the canonical MSC surface marker NT5E is repressed while inducing the adhesion receptor SORBS2 (Figure 3A; Mechanotransduction); none of these are induced or changed in the healthy donor. This suggests that transcriptional reorganization in HGPS adipogenesis is more primed towards a contractile cytoskeletal and increased adhesion state, which is opposite to rounding of cells during preadipocyte commitment, a plausible consequence of a lamina that cannot anchor cytoskeletal tension well.

#### Adipokines and fibrosis

SERPINF1, TNC, A2M, and LYVE1 are induced in both HGPS donors at both time points and in neither healthy donor at either time point. SERPINF1 is an anti-adipogenic adipokine associated with insulin resistance, and TNC and A2M are fibrosis markers, so the mechanical shift above is accompanied by a secreted fibro-adipokine signature (Figure 3A).

#### Alternative mesenchymal lineages

Consistent with this failure to transition out of the early, Wnt-inhibited state, both HGPS donors engage alternative mesenchymal programs despite specific adipogenic cues, but along different lineages. 11513 induces the chondrogenic ECM genes SOX9 [40], MATN3 [41], COL3A1 [42], COL5A1 [43] and MMP3 [44] are induced at D21 (Figure 3A and Supplementary Figure 3B), whereas 199 induces SOX9 and osteogenic marker BMP6 from D7, rising further by D21, without engaging the chondrogenic ECM set at all. Vascular smooth muscle programs are repressed in both donors, though at a different level of hierarchy: 11513 loses the regulators MYOCD [45] and NOTCH3 [46] (Supplementary Figure 3B), while 199 loses the structural genes MYH11 and DES. This pattern suggests partial leakage of mesenchymal fates during adipogenesis, which is also reported in the previous study from our lab in uninduced progeria donor fibroblasts [26].

#### Inflammation and SASP cytokines

These panels separate the two HGPS donors most sharply. In 11513, the classical SASP (Senescence-Associated Secretory Phenotype) cytokines IL1A and IL6 were repressed, while IL1RL1 is strongly induced at both timepoints; however, IL6 was also repressed in 08398 at D7, and IL1RL1 was induced in healthy at 1652 at D7, so only IL1A repression and sustained ILRL1 induction through D21 are specific to 11513. 199 instead represses SERPINB2 and induced IL1RL1 only weakly at D21 (Supplementary Figure 3B; SASP/inflammation). Engagement of the IL1RL1 receptor can target the downstream regulation of the JAK-STAT pathway, which plays a role in opposing adipogenic transcription [47].

Taken together, the HGPS donors show either discordant adipocyte markers at sub-healthy amplitude with chondrogenic drift (11513) or a more incompetent adipogenic program with stronger osteochondrogenic drift (199), yet both converge on a shared theme of mis-timed checkpoint engagement, altered mechanics, MSC identity loss, a fibro-adipokine switch and impaired lineage consolidation, a framework that helps explain lipodystrophy in HGPS.

### Adipogenesis is not uniformly blocked in HGPS donors: staging separates attenuated engagement from commitment without consolidation

To assess whether HGPS donors complete the canonical adipogenic differentiation program or stall at a particular maturation stage, we organized a set of 71 well-established adipogenesis markers into five sequential maturation stages: preadipocyte, early commitment, mid/lipogenic program, late terminal mature-adipocyte program, and lipolysis/mature-metabolism, and computed stage-mean z-scores together with a summary progression index (see Methods: Per stage RNA-seq and Hi-C chromatin stage trajectory analysis). The healthy 10-year-old donor 1652 displayed the canonical adipogenic trajectory: a monotonic rise of the late-terminal program across the three timepoints (Figure 3B), sustained positive mid-program engagement, and the highest terminal progression index of any donor. The healthy 8-year-old donor 08398 showed a quantitatively weaker but qualitatively similar pattern. Both progeria donors failed to recapitulate the healthy trajectory, but in distinct ways. 11513 (8-year-old HGPS) mounted a substantially smaller-amplitude transcriptional response than either healthy donor at every maturation stage; z-scoring across all donors, confirms that 11513 engages its own adipogenic program, but at an attenuated magnitude. The mid-program z-score was severely depressed at D7, and the late terminal-program z-score never crossed zero at either induced timepoint. In contrast, 199 (10-year-old HGPS) showed transient mid- and late-program engagement at D7 (late z-score comparable to healthy 1652 at the same timepoint) followed by regression at D21. Together, the staging analysis defines two distinct modes of HGPS adipogenic dysfunction, both distinct from complete failure: 11513 engages the adipogenic program at attenuated amplitude across all maturation stages. In contrast, 199 engages early and more broadly at D7 than 1652 but fails to consolidate, with FABP4 and LPL falling below 1652. Initiation is therefore mis-timed rather than delayed; individual markers move by D7, while coherent pathway enrichment emerges only at D21, and both models are comparable with the reported arrest in terminal maturation of HGPS adipogenic differentiation.

These distinct transcriptional failure models raise a fundamental question: are the adipogenic genes primed by or coupled to the underlying chromatin architecture of HGPS cells during adipogenic differentiation? HGPS is associated with the loss of spatial A/B compartmentalization and peripheral heterochromatin, along with altered epigenetic modifications [9, 48, 49], but which genomic regions are reorganized during adipogenic induction and how this reorganization contributes to the patient-specific lipodystrophy phenotype remains unclear. To address this, we used Hi-C data to first compare how MSCs and healthy fibroblasts reorganize their compartment landscape during adipogenesis, further comparing the healthy donor (08398) to HGPS donors (11513 and 199) to identify disease-specific architectural changes.

### Fibroblasts retain donor age and disease identity

We performed Hi-C experiments on fibroblasts from the 8-year-old healthy donor (08398) and from the 8-year-old (11513) and 10-year-old (199) HGPS patients. For each donor, Hi-C was performed at day 0 (D0, uninduced control) and day 21 (D21) under both control and adipogenic induction conditions. We compared these data with published bone marrow mesenchymal stem cell adipogenic differentiation Hi-C data [31]. We calculated the A/B compartment profile of each sample and then used Principal Component Analysis (PCA) to capture the variation in compartment profiles across all datasets. The results show that the 3D genome architecture separates cells based on disease state, donor, and differentiation state. (Figure 4B). PC1 (25.0% variance) places the MSCs as a discrete outgroup from the three fibroblast lines and, within each line, captures the shift from D0 to D21 induced state. PC2 (19.7% variance) stratifies samples by donor age: MSCs (youngest) at the bottom, 8-year-old healthy (08398) and progeria donor (11513) in the middle, and the 10-year-old progeria donor (199) at the top. Although 08398 and 11513 occupied the same PC2 band, they formed distinct D0 baseline clusters, indicating that the progerin phenotype is separable from donor age. These results suggest that donor-derived fibroblasts retain the biological genome structure variance of identity, age, and disease state. This contrasts with iPSC models, which have been widely used to study HGPS phenotypes, in which reprogramming and clonal selection reset donor age and disease-specific epigenetic architecture to a common pluripotent baseline [50, 51].

**Figure 4:**
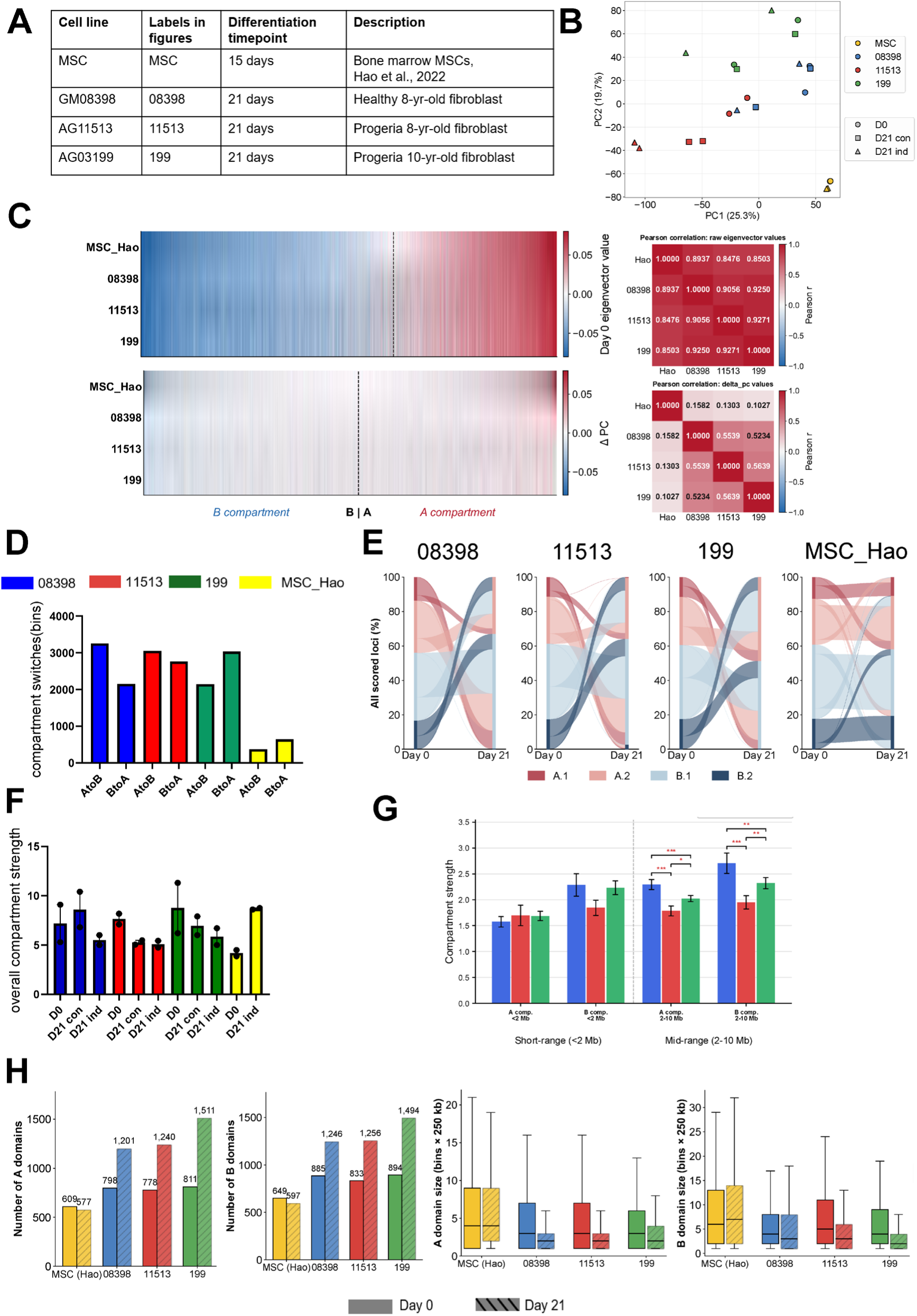
Adipogenic induction triggers widespread compartment switching in fibroblasts but minimal remodeling in MSCs. (A) Cell lines used for Hi-C, with figure labels, differentiation time points, and descriptions. (B) Principal Component Analysis of Hi-C compartment eigenvector profiles. Colors indicate cell lines, and shapes indicate conditions: D0, D21 (control), D21 (induced). (C) Top: Heatmap of D0 compartment eigenvector values across genomic bins, ordered by the MSC eigenvector. The dashed line marks the A/B boundary. Bottom: change in the eigenvector value between D0 and D21 induced condition (ΔPC), ordered by the MSC ΔPC. Matrices on the right show Pearson correlations between cell lines for D0 eigenvector values and ΔPC values. (D) Number of 250kb bins switching from A to B or from B to A between D0 and the D21 induced condition. (E) Alluvial plots show how loci move between sub-compartments from D0 to D21, as a percentage of all scored loci. For each cell line, A and B bins were split into strong sub-compartments (A.1 and B.2) and weak sub-compartments (A.2 and B.1). A bin was called strong if its PC1 value lay more than 0.5 standard deviations beyond the mean of that cell line’s own D0 A or B distribution. The same cutoffs were applied to the D21 data. (F) Overall compartment strength for each cell line and condition, calculated from saddle plots as (AA+BB)/(AB+BA). AA and BB are the mean observed/expected contact frequencies between the strongest 20% of A bins and the strongest 20% of B bins; AB and BA are the mean frequencies between A and B bins. Dots represent biological replicates; bars show the mean ± SEM. (G) Compartment strength of A and B compartments at short range (<2 Mb) and mid-range (2-10 Mb) in 08398 (blue), 11513(red), and 199 (green) at the D21-induced time point. Bars show the mean ± SEM across chromosomes. Statistical significance was assessed by the Wilcoxon signed-rank test, paired by chromosome (*p<0.05, **p<0.01, ***p<0.001). (H) Number of A and B domains (left) and their sizes (right) at D0 (solid) and D21 (hatched). A domain is a contiguous run of bins in the same compartment, and sizes are given in 250-kb bins. Boxes show the median and interquartile range.

### A shared baseline compartment landscape diverges during adipogenesis due to cell source and progerin presence

To more specifically compare cell lines by their 3D genome initial state and changes during differentiation, we ranked genomic bins by MSC D0 compartment eigenvector values and changes in these compartment values at D21 (ΔPC) for all cell lines, and calculated Pearson correlations between all compartment profiles (Figure 4C). At baseline, all cell lines share a highly similar bipartite compartment landscape. Correlation with the MSC reference is highest for the healthy (08398) fibroblast donor and lower for both progeria donors, while the two progeria lines are the most similar overall. Comparing the changes that occur during adipogenic induction, however, reveals more substantial differences between cell lines: ΔPC correlations between MSCs and fibroblasts fall to r=0.10-0.16, and among fibroblasts, they reach only 0.52-0.56. These results suggest that a shared bipartite compartment landscape is remodeled differently during adipogenesis across cell lines, with the remodeling response far less conserved than the baseline compartment identity. The divergence is structured rather than uniform: cell source separates more strongly, MSCs and fibroblasts share almost no common remodeling response, and disease state forms a weaker secondary grouping among fibroblast lines. Progerin is therefore associated with an altered trajectory of compartment reorganization in HGPS donor fibroblasts.

### Adipogenic induction collapses compartment strength in fibroblasts but strengthens it in MSCs

Comparing D21 to D0 timepoints, we observed extensive A↔B switching across all fibroblast lines (Figure 4D and 4E), compared with only 10% in MSCs. The fibroblast lines differed in directionality: 08398 was skewed towards compartment closing (A→B), whereas 11513 showed roughly equal numbers of opening and closing of compartments (Figure 4D). Both 199 and MSCs trended towards net compartment opening, with MSCs showing the strongest bias towards A, consistent with greater accessibility during lineage-specific genome organization [52, 53]. We further divided compartment switches into four category classifications: A.1= strong A, A.2=weak A, B.1=weak B, B.2=strong B, splitting each compartment at the mean ± 0.5 SD (standard deviations) of its own Day 0 PC1 subpopulation, with the same cutoff applied to the induction dataset to define sub-compartments. Beyond compartment switching, the alluvial plot revealed distinct fibroblast phenotypes (Figure 4E): A.1 and B.2 categories nearly disappeared, with virtually all D21-induced bins collapsing into the weak A.2 or B.1 categories. Hao MSCs retained substantially balanced strong and weak compartment populations on D21.

Genome-wide compartment strength dropped in all three fibroblast lines between D0 and D21 of the induction window, indicating that compartment weakening is an induction-specific response in fibroblasts. MSCs displayed the opposite trend: overall compartment strength increased across the same induction window (Figure 4F). Because compartment strength was calculated as genome-wide averages over all distances, we next asked whether the weakening of compartment strength was distance-dependent or uniform across scales during differentiation. To answer this question, we used the pentad tool [54] to stratify compartment strength by contact distance, the data indicated that short-range A/B segregation (< 2 Mb) was largely preserved across all fibroblast cell lines. In contrast, mid-range segregation (2-10Mb) was significantly reduced in the two HGPS lines during adipogenic lineage commitment (Figure 4G). This suggests that progerin-associated compartment weakening emerges primarily at the scale of compartment domains, indicating that segregation effects are more scale-selective than genome-wide.

### Fibroblasts fragment compartments into shorter blocks during adipogenic induction, closing gene-rich regions and opening gene-sparse ones

We next asked whether the adipogenic induction changed the granularity of the compartment landscape by grouping contiguous same-sign bins into “domains” (maximal run of A or B) and quantifying the number and size of these domains. All three fibroblast lines fragmented: compartments underwent striking fragmentation, resulting in more numerous and shorter same-compartment domains after adipogenesis induction, whereas the MSCs preserved both domain number and size (Figure 4H). The switches themselves showed the directional bias common to compartment-switch datasets, with closing (A→B) bins containing more gene-rich regions than opening (B→A) bins (Supplementary Figure 3C).

Together, these results indicate that MSCs and fibroblasts remodel their landscape towards adipogenesis differently. We therefore asked which chromatin signature distinguishes fibroblast adipogenesis from that of the MSCs and whether specific markers within this signature could identify the MSC -like phenotype within fibroblasts that enables adipogenesis, beyond the canonical MSC markers.

### A fibroblast-specific compartment signature identifies priming layers of adipogenic competence

We identified 420 genes unique to healthy adipogenesis in the context of fibroblasts (Supplementary Figure 4A) from the chromatin landscape (see methods: Novel adipogenesis chromatin signature in the context of fibroblasts). These genes fell within genomic regions that were in the A compartment in healthy fibroblasts and through adipogenesis, but which were in the B compartment in the ground state MSCs (Supplementary Figure 3D). GO terms organized these genes into five biologically relevant priming layers: (i) **Developmental TFs and Wnt signaling**, which is established to be an upstream regulatory cue for MSC lineage decision [39]; (ii) **Mitochondrial biogenesis and oxidative capacity** via Complex I subunits and mitoribosomal machinery, reflecting the mitochondrial expansion required for the energy demands of adipogenesis [55]; (iii) **membrane excitability and ion homeostasis** driven primarily by potassium transport, linking bioelectrical state to adipogenic priming [56]; (iv) **Niche cues via interleukin signaling**: the IL-4 receptor axis is engaged with its JAK/STAT effector, JAK1 [57]; and (v) **steroid and prostaglandin metabolism** via the aldo-ketoreductase family of AKR1C; this family is responsible for generating a PPARγ ligand (15d-PGJ_2_), a possible cue for activation of PPARγ during adipogenesis.

Having defined the chromatin priming architecture that supports healthy fibroblast adipogenesis, we next asked how this architecture is compromised in HGPS, focusing on the adipogenesis-relevant gene panel (see methods; Adipogenesis-focused concordance analysis). We identified compartment switches shared between both HGPS donors relative to the healthy 08398 donor. Further, we dissected the donor-specific features of 11513 and 199 that account for two distinct adipogenic failure models observed transcriptionally.

### A core HGPS signature of adipocyte closure and stalled adipogenic priming

We curated a 2213-gene panel for adipogenesis (see methods: Adipogenesis-focused concordance analysis); we mapped these genes to their respective 250kb Hi-C bins and plotted their ΔPC compartment changes on heatmaps shown in Figure 5 (see methods: Hi-C analysis). During adipogenesis, 154 genes consistently switch to B and 103 switch towards A in both HGPS donors, while these same genes move in the opposite direction in the healthy 08398 donor (Figure 5A).

**Figure 5:**
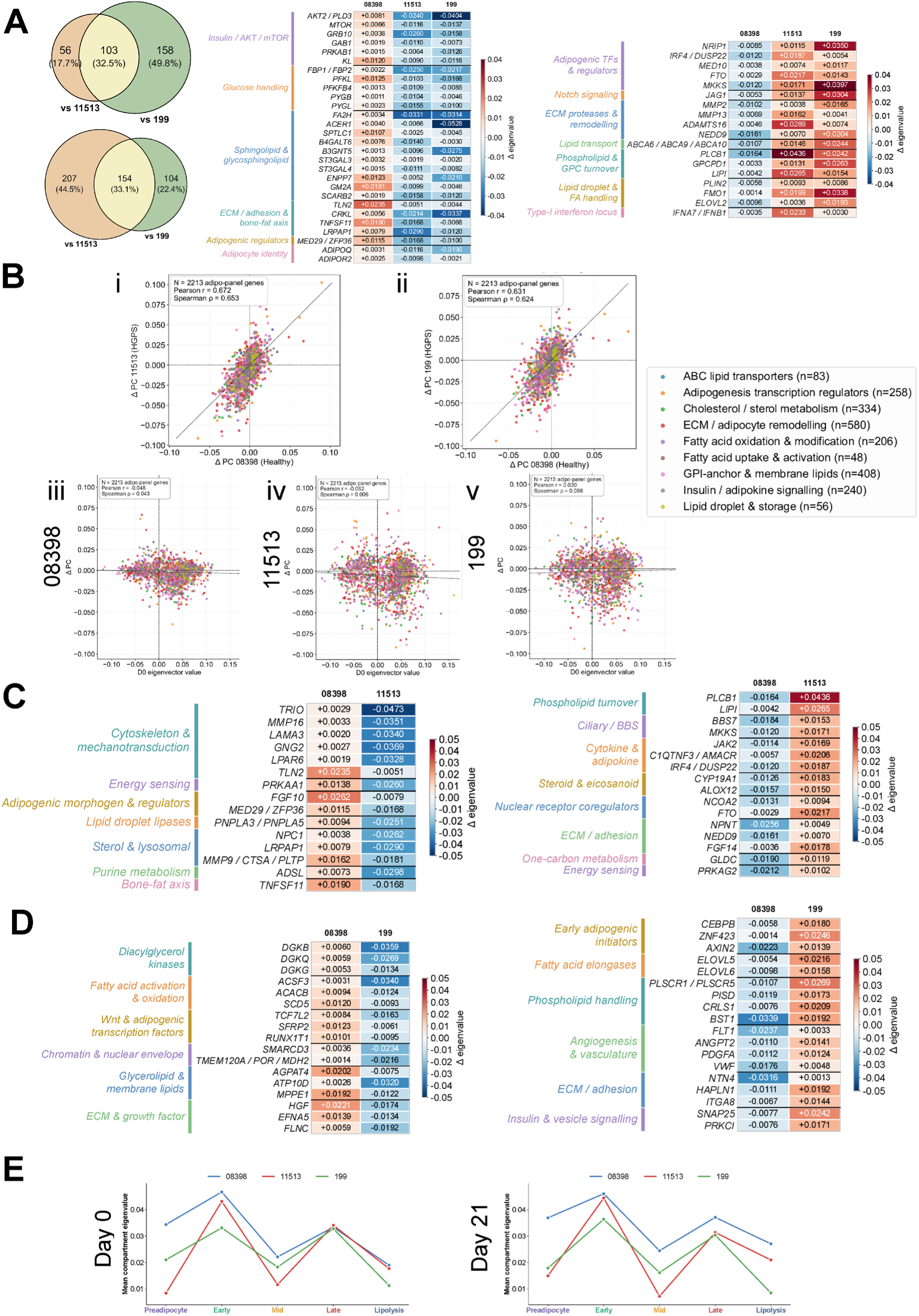
Progeria fibroblasts shift insulin and lipid metabolism genes towards the B compartment, and Notch, ECM, and interferon genes toward A, during adipogenesis. (A) Venn diagrams of adipogenic-panel genes whose change in ΔPC diverges between healthy (08398) and each progeria line (11513 or 199). The top diagram shows genes that gain A compartment in 08398 but lose it in progeria (shared n=103). The bottom diagram shows genes that lose the A compartment in 08398 but gain it in progeria (shared, n=154). The heatmap shows ΔPC values for selected shared genes, grouped by function: those gaining in 08398 and losing in both progeria lines (left), and those losing in 08398 and gaining both in progeria lines (right). (B) (i, ii) ΔPC of 2213 adipogenic panel genes in 08398 plotted against ΔPC in 11513 (i) and 199 (ii). Genes are colored by functional category. The dotted lines mark y=x. Pearson and Spearman correlation coefficients are shown. (iii-iv) ΔPC plotted against the D0 eigenvector values for 08398 (iii), 11513 (iv) and 199 (v). (C, D) ΔPC values for selected divergent genes specific to the 08398 vs 11513 comparison (C) or the 08398 vs 199 comparison (D), grouped by function. (E) Mean ΔPC values of the genes in each adipogenic stage module (preadipocyte, early, mid, late, and lipolysis) at D0 (left) and D21 (right) for 08398 (blue), 11513 (red), and 199 (green).

**A→B switches represent adipocyte program closure in the HGPS donors** (Figure 5A): The AKT2/PLD3 genes show the strongest ΔPC shift along with closure of mTOR, PRKAB1, GAB1, and GRB10, suggesting structural compromise of the insulin-AKT-mTOR axis, which could contribute to the lipodystrophy phenotype observed in the patients [58]. The glucose-metabolism loci genes like FBP1/2, PFKL, PFKFB4, PYGL also lose accessibility alongside insulin pathways. The sphingolipid pathway forms another major axis of closure in HGPS donors, predicting ceramide accumulation that could amplify insulin desensitization [59, 60]. In parallel, mature adipocyte identity genes such as **ADIPOQ** and **ADIPOR2** shift toward the B compartment [61]. This signature represents a coordinated chromatin closure for adipocyte identity, its insulin responsiveness, and metabolic substrate handling.

**B→A switches reveal a conflicting priming signature in HGPS patient-derived fibroblasts** (Figure 5A); we observe opening of pro-adipogenic transcription and anti-adipogenic cues simultaneously. Pro-adipogenic cues: HGPS donors move lipid droplet and fatty acid handling, phospholipid turnover, lipid transport channels, and ECM components for adipocyte remodeling into the A compartment. In parallel, the anti-adipogenic priming is co-occurring: PPARγ co-repressor *NRIP1 [62]*, the Notch ligand JAG1 (Notch signaling inhibits adipogenesis [63]) and type I interferon locus [64]. This co-opening of adipogenic initiators and anti-adipogenic cues suggests that HGPS cells engage the adipogenic program at the chromatin level but fail to fully resolve it, a state of “stalled” rather than committed priming in progeria cells.

Given their shared signature of impaired adipogenesis, we next asked how the two age-separated HGPS donors differ in the specific modes of dysregulation. To address this, we compared each HGPS donor directly to the healthy control donor, 08398.

### Global architecture is preserved but systematically displaced toward B in 11513

Comparing ΔPC values across all 2213 adipogenic panel genes, healthy (08398) and HGPS donor (11513) are correlated, suggesting that broad adipogenic compartment architecture is retained in 11513 (Figure 5B, i). Within this retained subset of genes in 11513, most of them move towards B in comparison to 08398, consistent with the asymmetric switching observed in this sample (A→B = 361; B→A = 159). In 11513, the mean displacement of all 9 functional categories in the adipogenic panel moved toward B, and none moved toward A on average, although genes within individual categories do. The categories differ only in magnitude: displacement is the largest for insulin/adipokine signaling and cholesterol metabolism, followed by lipid droplet and storage.

The shift into the repressive B compartment is global, but greatest for insulin-responsive genes, sterol handling, and lipid storage: three functions most directly required to build and maintain a mature adipocyte.

To identify loci within these functional categories that may drive the dysregulation in 11513, we prioritized genes with the strongest B-ward shift within each category. Notably, the most pronounced shifts fall at cytoskeletal and adhesion loci rather than metabolic genes, a pattern that points towards the lamina defect itself rather than its metabolic consequence (Figure 5C, left). TRIO shows the largest shift, alongside MMP16, LAMA3, TLN2, LPAR6 and GNG2, a cohort that builds the cytoskeleton and anchors the cell to its matrix. Because the nuclear lamina anchors the LINC complex, the physical link through which cytoskeletal tension reaches the genome, a Lamin A/C defect would therefore be expected to disrupt this connection, and compartment reorganization at these may be its chromatin-level readout. Alongside the mechanical cues moving towards the B compartment, we also observe closure of fat cell identity loci FGF10 [65] and PNPLA3/PNPLA5 lipase modules that are important for lipid droplet biogenesis [66].

Fewer genes move into the open compartment, and they fall into several groups. PLCB1 is the largest shift, along with LIPI; both act in phospholipid turnover and are in the A compartment in 11513 (Figure 5C, right). Two Bardet-Biedl syndrome genes, BBS7 and MKKS, become accessible; notable because primary cilia are required before a precursor cell can become an adipocyte [67, 68]. PPARγ co-activator NCOA2 [69] and pro-adipogenic factor FTO open [70], along with opening of inflammatory signals, which are anti-adipogenic cues, suggesting that activating and opposing machinery gain accessibility together in the HGPS donor.

### Balanced bidirectional remodeling in 199 primes matrix remodeling at the expense of adipocyte maturity

Across all adipogenesis panel genes, 199 strongly correlates with 08398, similar to 11513; therefore, the broad organization of the adipogenic panel is retained (Figure 5B, ii). However, the direction of change in 199 is bidirectional and presents a symmetric switching pattern, unlike 11513 (A→B=258, B→A=261). No functional category is directionally distinct in 199; even within each functional class, there is more duality than a uniform opening or closing of compartments.

The individual loci that close in 199 belongs to the lipid-handling machinery (Figure 5D; left). The three diacylglycerol (DAG) kinases move together (DGKB, DGKQ, DGKG); they are responsible for converting diacylglycerol to phosphatidic acid. Without the conversion, accumulation of DAGs can lead to insulin resistance and can potentially lead to impaired insulin signaling [71]. In parallel, fatty acid oxidation and activation loci also close (ACSF3, ACACB, SCD5, ACSM4, ACOT13) along with glycolipid, cholesterol, and membrane lipid loci.

Apart from the metabolic block that the 199 HGPS donor undergoes, we observe (i) Wnt signaling and adipogenic loci closing together, like TCF7L2, a type-2 diabetes risk locus [72], SFRP2, and RUNX1T1. (ii) SMARCD3/BAF60c, which is a chromatin remodeler responsible for specific lineage commitment [73]; its closing may provide a chromatin-level correlate for the pronounced mesenchymal leaky expression we observe by RNA-seq. (iii) TMEM120A/TACAN, a nuclear envelope protein essential for fat cell differentiation [74].

199 donors have experienced shifts to open compartment identity for early adipogenic initiators like CEBPB, ZNF423, and AXIN2, unlike 11513 (Figure 5D; right). We also observe fatty acid elongases and phospholipids in compartment A. This opening of lipid breakdown machinery, alongside the shutdown of lipid-building loci described above, suggests an unresolved state in the commitment program that 199 is engaged in. Active remodeling during adipogenesis is a notable feature of 199, with matrix components and proteases poised in the A compartment. Therefore, 199 elaborates a signature where adipogenic initiation and matrix component loci are open but is closed for lipid and metabolic effectors.

### HGPS fibroblasts begin from a shared chromatin baseline but reorganize chaotically during differentiation

Having identified the loci that switch in shared HGPS donor and individual donor, we next asked whether these differences are present at the baseline or emerge over the differentiation window. On D0, the compartment architecture was highly concordant across donors for all 2213 adipogenic panel genes, indicating that healthy and HGPS fibroblasts have the loci primed at a common chromatin configuration (Figure 5B, iii, iv, v). During induction, 08398 changes compartments at a modest magnitude or constrained dispersion, whereas HGPS donors showed a threefold increase in the dispersion of the loci being remodeled. Remodeling in HGPS is therefore more variable from locus to locus than in the healthy donor, rather than uniformly larger in one direction. Also, the two patient lines are disordered in different ways; in 11513, the initial A compartment loci lost compartment strength more often than they gained, consistent with more A→B bin switches (Figure 5A). In 08398 and 199, there was a balance to the switching, suggesting directional bias as a donor-specific trend in 11513.

Baseline compartment strength did not predict differentiation-induced change in any cell line; correlations between D0 and ΔPC were negligible and unstable. We therefore do not interpret D0 values as predicting how far a locus subsequently moves during differentiation.

### HGPS fibroblasts undergo genome-wide compartment reorganization during induction without achieving stage-specific compartment identity

To understand at which stage is the adipogenic induction failing at the chromatin level in HGPS donors, we assembled a 71-marker panel spanning 5 sequential stages of adipocyte (see methods: Per stage RNA-seq and Hi-C chromatin stage trajectory analysis): Preadipocyte, Early, Mid/lipogenic, Later/terminal, and Lipolysis/metabolism. Mean compartment eigenvector values were plotted across D0 and D21 induced time points (Figure 5E). At D0, 08398 shows the most strongly primed chromatin landscape for pre- and early-adipocyte markers, whereas both HGPS donors fall below this trajectory, consistent with reduced priming of mid-stage/lipolysis-associated loci before induction. After induction, 08398 retains its pre/early marker configuration but shows increased engagement of mid-to lipolysis-stage loci, suggesting that induction preferentially remodels this mid-stage to lipolysis module in healthy fibroblasts. In contrast, both HGPS donors fail to engage the mid-stage module and show only modest late-stage remodeling relative to D0. This chromatin-level pattern mirrors the transcriptomic trajectory, indicating that the RNA-level failure is also anchored into the donor-specific compartment architecture. Notably, although 11513 shows increased compartment engagement at lipolysis stage loci, this does not appear to translate effectively at lipid droplet maintenance [75], consistent with the reduced BODIPY+ lipid droplet size we observed by imaging (Supplementary Figure 1B).

Having established that chromatin and transcriptomic trajectories are broadly aligned, we next asked whether this concordance extends to locus-level coupling between compartment status and gene expression. To address this, we related per-gene A/B compartment shifts to per-gene transcriptional changes across the induction window.

### A/B compartment identity is only partially coupled to transcriptional output during adipogenic differentiation

To assess the temporal relationship between compartment and transcription, we compared the log2fold change at D7 and D21 against D21 compartment Δpc, per gene. Concordance percentages were similar across RNA time points-66% in 08398, ∼60% in 11513, and 59% in 199, indicating the compartment-transcription coupling is broadly established by D7 and extended through day 21 (Figure 6A, left). Individual genes behave the same way: every gene that is differentially expressed at both times retains the same concordance of directionality (Figure 6A, right). The D21 compartment state is therefore equally concordant with early and late transcriptional change; because compartments were profiled at D0 and D21 only, we treat this concordance as concurrent rather than as an ordering of the two.

**Figure 6:**
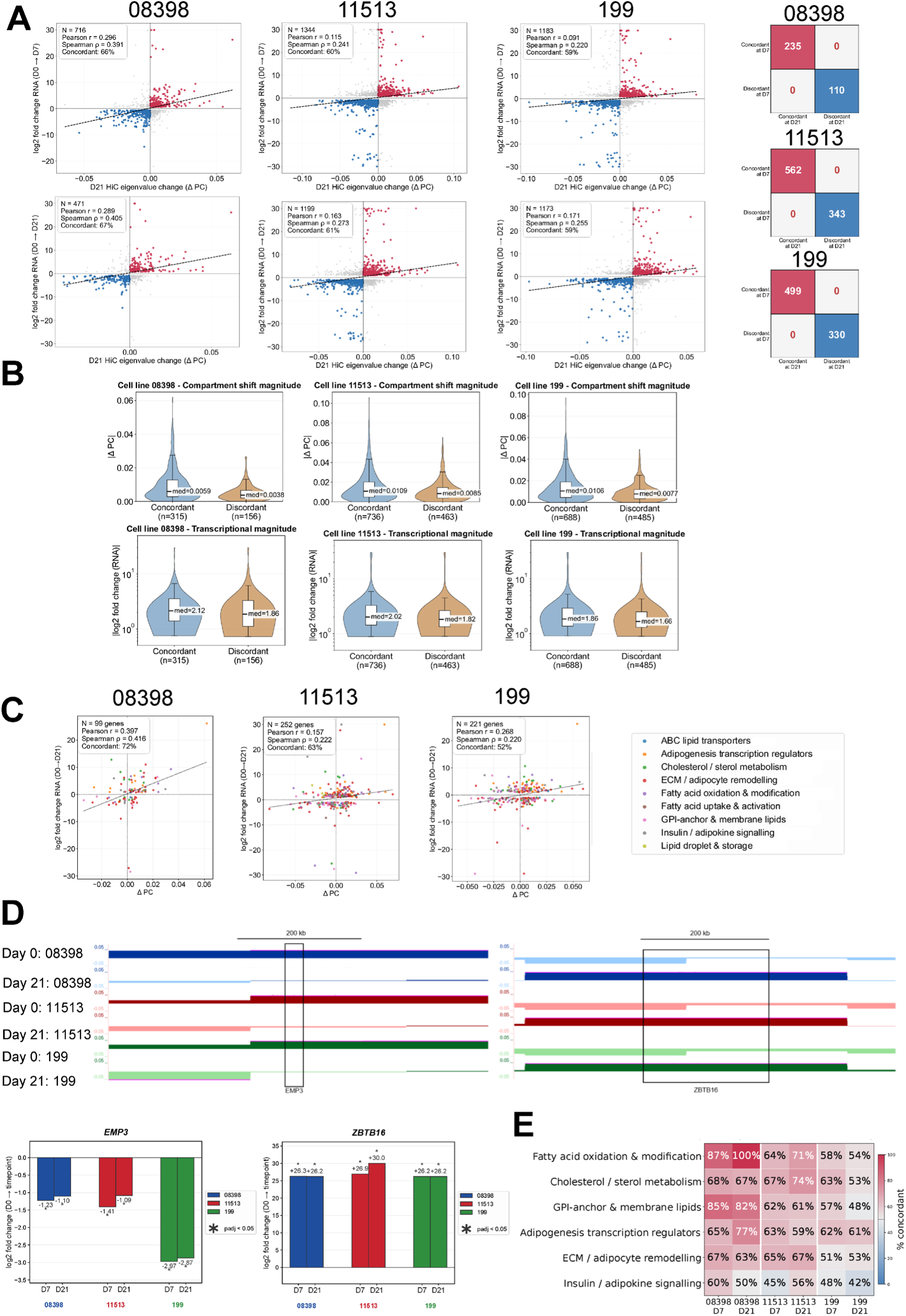
Compartment changes are less tightly coupled to gene expression in progeria fibroblasts. (A) Log2fold change in RNA expression (D0→D7, top; D0→D21, bottom) plotted against the change in Hi-C ΔPC for D0 and D21 for each gene in 08398, 11513, and 199. Red and blue points are DEGs with positive and negative values, respectively, for genes that change expression and Hi-C compartment identity in the same direction. Dashed lines show the linear fits. The number of genes, Pearson and Spearman correlation coefficients, and percentage of concordant genes are shown in the plot. The tables on the right compare each gene’s concordance status at D7 with its status at D21. (B) Absolute ΔPC (top) and absolute RNA log2fold change at D21 (bottom, log scale) for concordant and discordant genes in each cell line. Medians are shown. (C) RNA log2fold change (D0→D21) plotted against ΔPC for adipogenic panels, colored by functional categories. (D) genome browser views of EMP3 (chr19) and ZBTB16 (chr11) loci (hg19). For each cell line, the tracks show D0 eigenvector values (light) and ΔPC (dark); 08398 in blue, 11513 in red, and 199 in green. The bar plot shows RNA log2fold change at D7 and D21 (*padj<0.05).

Having established that 60-66% of DEGs at D21 move in the compartment-predicted direction, we asked what distinguishes the concordant and discordant cohorts: is it the magnitude of transcriptional response or the magnitude of their local compartment shift, or both? For every cell line, we compared the log2fold change in RNA and Δpc compartment values between the two cohorts on D21. For both cohorts, transcriptional magnitude overlapped completely; however, the compartment shift magnitudes differed dramatically. The concordant cohort occupied bins that underwent larger compartment organization, while the discordant cohort occupied bins with weaker magnitude of compartment shifts (Figure 6B). The results indicate that A/B compartment identity is only partially coupled to transcriptional output, and this coupling operates in a permissive rather than instructive mode: strong compartment reorganization is largely accompanied by concordant transcriptional change, but modest compartment reorganization is compatible with transcriptional decoupling in either direction. Figure 6D shows representative genes with concordant compartment and expression changes across all three cell lines, displayed as compartment tracks alongside log2 fold-change values.

### Chromatin-transcription coupling weakens with donor severity and collapses first in lipid-metabolic pathways

To examine which biological programs of adipogenesis are most tightly aligned with the D21 compartment landscape and how this alignment evolves during induction, we computed the concordant (RNA-compartment-aligned) genes per adipogenesis panel and per cell line at D7 (Supplementary Figure 4B) and D21 (Figure 6C) (see methods for details).

Coupling was strongest in 08398 (healthy) and declined across two HGPS donors on D21. The same trend holds when D7 expression was compared to the D21 compartment landscape, indicating the relationship is not an artifact of measuring RNA and Hi-C simultaneously. Notably, this loss of coupling in HGPS donors cannot be attributed to a weaker transcriptional response: HGPS patients have twice as many DE genes represented in the adipogenic panel as 08398 at D21. HGPS cells therefore mount a transcriptional response of similar magnitude and greater breadth during adipogenic induction but are progressively uncoupled from the underlying compartment landscape.

To identify which functional programs dominate the loss of coupling in HGPS, we resolved RNA concordance by adipogenic panel category (Figure 6E). In 08398, the lipid-metabolic arm is the most tightly coupled: fatty acid oxidation reached 87% at day 7 and 100% by day 21, followed by the GPI anchor/membrane lipid biosynthesis category. Both categories collapse in HGPS donors; cholesterol/sterols metabolism alongside ECM remodeling are the more stable categories in HGPS donors, but collapse progressively with donor age, as seen in 199. The only adipogenic panel program that was preserved in 199 is the transcription regulators in comparison to the healthy. Insulin/adipokine signaling is the weakest program across all cell lines and timepoints, suggesting that this arm is uncoupled from the compartment state irrespective of the donor.

Together, these results indicate that compartment position and transcription are partially coupled in healthy adipogenesis, and that this coupling is progressively lost in HGPS. The decoupling is not uniform across the adipogenic program: lipid-metabolic categories are tightly coupled in healthy fibroblasts but are the first to decouple in HGPS donors; these programs can contribute to a compromised adipocyte identity that manifests at the tissue-level biology of HGPS patients.

## Discussion

Taken together, our results show that HGPS patient-derived fibroblast cultures commit to the adipogenic lineage at frequencies comparable to age-matched controls but fail to stabilize that commitment, retaining access to competing mesenchymal programs and producing morphologically incomplete adipocytes. This failure is accompanied by a scale-selective loss of mid-range compartmentalization, a threefold increase in the variance of compartment remodeling, and progressive uncoupling of the lipid-metabolic transcriptional program from the underlying chromatin architecture.

### Consolidation, not induction, is the point of failure

Initiating a lineage program by signal transduction and master regulators is different from consolidating the lineage program; this is a slower process by which the cell restricts remaining options, stabilizes the new program, and builds a structure that is crucial for the differentiated cell state. Our data place the HGPS defect in the latter of these.

The commitment outcome is reached in HGPS patient-derived cells, as shown by a similar frequency of FABP4+ cells as in the healthy age-matched fibroblast cultures, the induction of PPARγ effectors like LPL, CIDEC, LEP, PLIN1, and the accumulation of lipid droplets. However, we observe failure in maturation across several independent readouts. Lipid droplets form but stay smaller in HGPS lines in comparison to WT controls, cage like structures around the lipid droplet are heterogeneous or absent in HGPS cells, and CEBPA gene expression induction is absent in both HGPS donors while reaching significance in both healthy donors. These are all consolidation events rather than entry events for adipogenesis; CEBPA in particular is dispensable for commitment but required for functional competence [76]. Therefore, HGPS cells reach the adipocyte state and stall within it.

Our data also suggests that the lineage restriction also fails in parallel. HGPS donor fibroblasts subjected to adipogenic induction show roughly twice as many DEGs as healthy fibroblasts, out of which 57-62% are absent from either control at any time point. Compensation for a weakened induction could explain elevated output but not its composition: SOX9 and BMP6 are transiently induced in wild-type cells at D7 but resolve by D21, whereas both these gene signatures persist and amplify by D21 in HGPS donors. This indicates that HGPS holds alternative lineage commitment options open throughout adipogenic induction.

We also observe that the failure is not redirected; DEGs shared between healthy donors are rarely discordant in progeria donors, and DEGs expressed at both time points mostly maintain their directional regulation across donors. Therefore, this suggests that the program initiates correctly but remains unsupported through adipogenic progression.

What sustains an initiated adipogenic program? We propose that genome organization supplies part of this support, selectively rather than uniformly. Only two-thirds of DEGs change gene expression in the direction predicted by their shift in spatial compartment. However, certain pathways are much more strongly coupled to compartment changes than others. In induced healthy fibroblast cultures, fatty acid oxidation DEGs show complete concordance with compartment changes, while insulin and adipokine signaling pathways are only weakly coupled to compartment shifts throughout progression. This suggests that genomic architecture underwrites the lipid-metabolic arm. Notably, it is this set of pathways that become misregulated in HGPS, and discordance between gene regulation and compartment change is more likely in these genes during adipogenesis.

### The coordination of adipogenic cues is defective at the genomic architectural level

If genome organization underwrites the programs that fail in HGPS, when does the architectural defect itself appear? Prior work describes progressive compartment weakening with passage, with TAD structure retained [26, 48]. Our data point to a failure of coordination rather than a particular compartment state. At early passages, baseline architecture is fairly concordant between healthy and HGPS donors, in contrast to the compartment changes reported in the later-passage HGPS fibroblasts [48] but increased divergence emerges with adipogenic induction, as a threefold increase in dispersion of compartment shifts occurs with no directional bias. Therefore, HGPS cells diverge from healthy controls in largely line-specific directions; progerin removes the constraint that makes the remodeling and reorganization during adipogenesis coherent and reproducible.

### The scale of the defect implicates the lamina

Our distance-stratified genome architecture analysis discriminates among the ways progerin could compromise genome organization. We find that short-range A/B segregation below 2Mb is preserved in every fibroblast line, while segregation between 2-10 Mb is weakened in both HGPS donors. This is the length scale of Lamina-associated domains and compartment domains rather than of cohesin- and CTCF-dependent loops and is consistent with reports that TAD structure is intact and retained [26, 48]. This scale of disruption is consistent with loss of peripheral heterochromatin and altered H3K9me3 and H3K27me3 landscapes described in HGPS [8, 9, 77].

### Adipogenesis proceeds by more than one architectural route; the fibroblast route is lamina-dependent

By comparing MSC and fibroblast adipogenic induction, we observed opposite architectural responses in the two cell types: MSCs sharpen compartment segregation and increase cross-compartment mixing, while fibroblasts weaken compartment strength and fragment domains. Since both experiments yield adipocytes, we propose that the route is set by the starting configuration: a MSC selects amongst latent lineages whereas a fibroblast must dismantle an established somatic identity. These remodeling tasks act on the compartment landscape, and the lower adipogenic efficiency of fibroblasts is consistent with dismantling being harder of the two. Why the fibroblast route is harder is not obvious from the compartment data alone, but it has a mechanical precedent: Lamin A scales with tissue stiffness; adipogenesis is favored by low Lamin A, and higher Lamin A impedes nuclear remodeling under stress [78]. Fibroblasts form a stiffer matrix than uncommitted progenitors and thus begin in a remodeling-resistant configuration.

### Reconciling with previous reports of impaired commitment

Our finding that FABP4+ frequencies after adipogenic induction are comparable between healthy and HGPS donors appears to conflict with reports of impaired adipogenic potential in HGPS. Xiong and colleagues [22] described progerin-dependent repression of PPARγ and C/EBPα induction network in iPSC-derived cells, and Najdi and colleagues [23] reported reduced adipogenic capacity in skin-derived precursors. Two considerations may reconcile these discrepancies: (i) the starting population differs in competence: both previous studies used stem or progenitor populations in which most cells are adipogenically competent, so reduced differentiation reflects a per-cell deficit. In our heterogeneous fibroblast cultures, only a small subpopulation is competent in either healthy or HGPS donors, so FABP4+ frequency reports the size of that subpopulation along with the potential capacity of each cell. (ii) the readout captures different quantities, commitment frequency versus maturation quality, which our morphological data show diverge. Notably, our observations showing that C/EBPα induction fails while PPARγ effectors are induced is consistent rather than conflicting with the model proposed by Xiong and colleagues, differing in which node of that network is affected.

### Implications

If the HGPS adipogenesis defect is consolidation rather than induction, the therapeutic objective is not to drive more cells into the adipogenic lineage but to allow the committed cells to complete and retain maturation. Several interventions follow, each with a defined readout in this model. Exogenous PPARγ ligand would test whether ligand availability limits maturation, since the receptor expression is intact and its effectors are induced while the downstream remodeling effector FGF1 is repressed in both HGPS donors. JAK-STAT inhibition addresses the type I interferon locus that gains accessibility in both donors and is the only candidate with direct support in this disease. Baricitinib, a JAK inhibitor, has been previously shown to improves adipocyte differentiation and lipid droplet formation in skin-derived precursors from HGPS and two other laminopathies, both alone and combined with farnesyltransferase inhibitor [79], in the same system in which reduced adipogenic capacity was originally reported [23]. Notch inhibition targets JAG1, which is shared by both donors. Notch signaling maintains mesenchymal progenitors in an undifferentiated state [63]; γ-secretase inhibition raises CEBPA in these cells [80], the single induction event absent in both HGPS donors and present in both controls, so the pharmacology and the defect meet at the same gene. Because all three interventions act downstream of progerin rather than on it, each could complement rather than replace progerin-targeted therapy.

Evaluating any of such interventions in patients would require a readout that depot mass does not provide. Farnesyltransferase inhibition improves rates of weight gain in a subset of patients, and whole-body DXA attributed that gain to fat at lower weight-gain thresholds and to lean mass at higher ones, with the authors noting limited sensitivity of the method [81]. Adipose mass is in any case an indirect measure of the process described here, since a depot can expand through hypertrophy of existing adipocytes without any cell completing maturation. Consistent with this, fat mass index is normal in HGPS children while trunk-to-leg and android-to-gynoid ratios are elevated [82], suggesting an adipose phenotype of distribution and organization rather than absolute depletion. Adipocyte number and size distribution, which our model identifies as the informative measurements, have not been assessed in patients.

More broadly, a defect expressed as variance rather than as a mean shift is invisible to differential analyses that test for consistent change, which may bear on how architectural phenotypes are sought in other disease and aging contexts.

## Conclusion

In this study, we conclude that progerin does not close the door on the adipogenic lineage. It removes the constraint that normally makes differentiation reproducible, leaving cells that commit correctly, transcribe extensively, and cannot finish. Lipodystrophy in HGPS is therefore better understood as a failure of consolidation than as a failure of commitment, and the architectural defect underlying it is a loss of coordination rather than the adoption of a wrong configuration.

The impact of this work is threefold. Relocating the lesion from commitment to consolidation changes what a therapeutic intervention must achieve and what a successful one would look like. Recognizing that the architectural phenotype resides in the spread of compartment change rather than in its direction identifies a class of defect that conventional differential testing, which compares group means, is not designed to detect. And showing that compartment-level changes can nominate therapeutic targets that expression data alone do not surface offers a route from genome organization to pharmacology. The object for correction is the capacity to remodel, not the expression of any individual gene.

## Methods

### Cell lines and culture

Age-matched healthy and progeria fibroblasts were obtained from the Coriell Institute for Medical Research: healthy lines GM08398 (8-year-old) and GM01652 (10-year-old) were cultured in MEM (Gibco, Cat. #12561072) supplemented with 15% FBS, 1X penicillin-streptomycin, and 1X L-glutamine and 1X non-essential amino acid (Gibco Cat. #11140050) at 37°C, supplied with 5% CO_2._ Hutchinson-Gilford Progeria Syndrome lines AG11513 (8-year-old) and AG03199 (10-year-old) were cultured in Rooster Nourish MSC media (Cat. #K82003) to increase the yield and viability of the fibroblasts. After expansion, the cells were transferred to the normal culture medium: MEM (Gibco, Cat. #12561072) supplemented with 15% FBS, 1X penicillin-streptomycin, and 1X L-glutamine. All experimental data were collected after 3-4 passages of acclimatization to the normal medium.

Human bone marrow-derived mesenchymal stem cells (hMSC-BM; PromoCell) were obtained from the manufacturer and cultured according to the recommended protocol. Briefly, cells were maintained in a Mesenchymal Stem Cell Growth Medium 2 (PromoCell Cat. #C-28017), prepared by mixing the basal medium with the provided Supplement Mix. The cells were seeded into T75 flasks at the manufacturer-recommended initial density of 4000 cell/cm^2^ and cultured in a humidified incubator at 37°C supplied with 5% CO_2_. Complete growth medium replacement was performed every 2 to 3 days, and cell growth was monitored regularly until a confluency of 70– 90% was achieved.

### Adipogenic differentiation for imaging and RNA collection

Sterile German-glass round coverslips (Cat. #76305-514) were placed in a 24-well plate, and 50,000 cells were seeded per well in MEM Alpha (1X; Gibco, Cat. #12561-056). At confluence, adipogenic differentiation was induced using StemXVivo Adipogenic Supplement (R&D Systems, Cat. #CCM011; 3.5 mL) and carried out for 7 and 21 days.

For imaging, cells were incubated overnight with BODIPY (Invitrogen, Cat #B3932), diluted 1:500 from a 1mg/mL stock concentration. Cells were then fixed in 4% PFA and stained for FABP4 (R&D Systems, Cat. #AF1443) according to the manufacturer’s protocol (R&D Systems, Cat. #SC006) followed by incubation with an Alexa Fluor 488-cojugated donkey anti-goat secondary antibody (Invitrogen, Cat # A-11055). Nuclei were counterstained with DAPI using either mounting (Invitrogen, Cat # P36966) or by incubation with DAPI diluted from 1:1000 a 1mg/mL stock (Invitrogen, Cat #62248). Images were acquired on an EVOS FL Auto II imaging system with an Olympus Plan Apo 40x objective and GFP and Cy5 fluorescent light cubes. FABP4+ cells and morphology were quantified manually per field.

For RNA isolation, cells were lysed in 50 µL RLT buffer per well, and RNA was extracted using the RNeasy Micro Kit (Qiagen, Cat. #74004). RNA quality and quantity were assessed by NanoDrop. Total RNA was submitted to GENEWIZ for poly(A) mRNA library preparation and sequencing.

### RNA-seq analysis and shrunken log fold change

All the RNA-seq reads were first processed with the BBDuk tool (https://github.com/kbaseapps/BBTools). Adapter trimming was performed, followed by quality trimming to discard reads below a quality score of 28. The reads were then aligned to the hg19 reference genome with STAR 2.7.6a (https://github.com/alexdobin/STAR). HTSeq 0.11.2 (https://github.com/htseq/htseq) was then used to obtain feature counts. To prepare for differential gene expression analysis, features with less than 10 counts across all samples were filtered out. Since the RNA-seq files were generated in three separate experiments, the raw gene count files suffer from batch effects. ComBat-seq (https://github.com/zhangyuqing/ComBat-seq) was used to adjust the data and account for potential batch effects. Files produced in the same experiment run are assigned the same batch number. Additionally, the sample origin and collection day are provided as biological covariates to preserve that signal in the final adjusted data. DESeq2 (https://bioconductor.org/packages/release/bioc/html/DESeq2.html) was used to determine differential gene expression between samples and/or conditions. The resulting data was then run through the lfcShrink function using the apeglm (https://bioconductor.org/packages/release/bioc/html/apeglm.html) estimator to shrink log2 fold changes for lowly expressed or noisy genes.

### MSC vs fibroblast transcriptional profiling

To characterize stem cell marker expression across MSCs, healthy fibroblasts, and HGPS donors, we curated a stem cell marker set from five Enrichr libraries (MSigDB_Hallmark_2020, KEGG_2021, Reactome_2022, GO Biological Process_2023 and WikiPathway_2023_Human) and intersected it with our RNA-seq data. Pathways were filtered by keywords to retain stemness-relevant terms and exclude EMT or cancer-associated pathways. Each gene was assigned to one functional category by its strongest pathway overlap. ComBat-seq counts were converted to log_2_(TPM+1). Of 1136 markers detected in our RNA-seq data, 505 have concordant replicates, defined as |R1-R2|≤ 1.5 log_2_(TPM+1)in every one of the 22 conditions. Markers were grouped by expression pattern using a> 2-fold difference in mean expression between cell groups: MSC restricted, HGPS lost, healthy fibroblast-specific, or induction impaired.

### Pathway-level log2FC plot

Genes for each pathway category were identified by GO Biological Process enrichment analysis Enrichr [83–85] and manually curated functional panels spanning compromised pathways reported in the HGPS literature, like Wnt signaling [32–34], cell cycle [34–36], oxidative stress, skeletal and vascular programs, and adipocyte markers [10]. Genes from these panels were plotted if they were differentially expressed (p adj <0.05) in at least one donor at D7 or D21, and each HGPS donor is shown alongside its age-matched healthy control so that HGPS-specific, shared, and healthy specific changes can be compared.

### Adipogenic differentiation assay for Hi-C experiments

Cells were seeded onto glass untreated coverslips in a 6-well plate and subjected to adipogenic differentiation for 21 days using the reagents described above. Nuclei were isolated prior to Hi-C, and pellets were fixed using the low-input protocol, followed by the Arima-HiC+ Kit protocol for mammalian cells (Arima A160134). For the low-input fixing protocol, the cells were washed with 3% BSA and crosslinked with 2% formaldehyde for 10 minutes at room temperature, then add 91.9 µL of stop solution 1 and incubate for 5 minutes at room temperature, then incubate on ice for 15 minutes, wash it with 3% BSA and snap freeze and store at -80°C or proceed with Arima Hi-C protocol. Libraries were prepared with the Arima Library Prep Kit, set 1 (Indexes 1–16), with PCR amplification cycles optimized for low-input cell counts based on sample yield. Sequencing was performed at GENEWIZ using paired-end 150 bp reads.

### Hi-C analysis

Sequencing reads were mapped to the human genome (hg19), filtered, and iteratively corrected by HiCPro (https://github.com/nservant/HiC-Pro). Library quality and mapping statistics are presented in Supplementary Table 1.

Compartment analysis was performed by running principal component analysis on 250 kb binned matrices using the matrix2compartment.pl script in the cworld-dekker pipeline available on GitHub (https://github.com/dekkerlab/cworld-dekker). The PC1 value was then used to represent the compartment identity for 250 kb bins.

PCA on these compartment level signatures across samples revealed two competing sources of variance: differentiation state (D0, D21 control, D21 induced) and variation from passage number and replicate-specific drift. Direct comparison of raw PC1 values between any two samples would therefore conflate the adipogenic signal with passage-acquired differences. To isolate the differentiation-attributable component, we adopted a per-bin replicate-averaging strategy: for each 250 kb bin in each cell line, raw PC1 values were averaged across the two biological replicates separately for each condition, baseline-corrected against D0, and the induced-minus-control differential retained as the final per-bin signal:

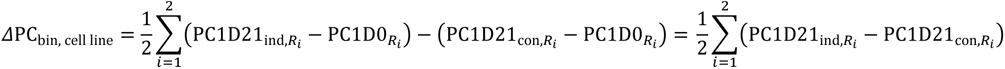

If the two replicate ΔPC values had opposing signs, the bin was excluded from further analysis rather than averaged. This removes replicate-specific scatter, passage-related drift, and changes due to time in culture shared between control and induced arms, isolating the chromatin reorganization specifically attributable to adipogenic induction.

Gene-level ΔPC computation from Hi-C bins

To translate bin-level compartment eigenvalue changes into gene-level values, the 250-kb bin table was collapsed to a per-gene, per-cell-line ΔPC (as calculated above) using GENCODEv19. Bins with missing gene annotation or missing ΔPC were skipped. For genes spanning multiple bins, each gene’s cell-line-specific ΔPC was computed as the arithmetic mean of ΔPC across all bins.

### Adipogenesis-focused concordance analysis

An adipogenesis gene panel was assembled by pathway-enrichment lookup. Full gene-set libraries from five Enrichr databases were used: MSigDB Hallmark 2020, KEGG 2021 (human), Reactome 2022, GO Biological Process 2023, and WikiPathways 2023 (human). Nine functional categories were defined prior: fatty acid uptake/activation; fatty acid oxidation/modification; lipid droplet and storage; cholesterol and sterol metabolism; ABC-family lipid transporters; adipogenic transcription regulators; insulin and adipokine signalling; extracellular matrix and adipocyte remodelling; and GPI-anchor and membrane lipid biosynthesis. Each category was defined by 5– 11 keyword strings (e.g. ‘ppar signal’, ‘β-oxidation’, ‘lipid droplet’, ‘gpi’, ‘adipokine’). Pathways whose names matched any keyword were retained, and the union of their genes was intersected with the compartment dataset (∼51,000 genes with non-zero mean ΔPC in ≥1 cell line). Each gene was uniquely assigned to the category whose matched pathways contained it most frequently.

### Saddle Plots

Multiresolution cool files were created from the HiCPro validpair files using the cooler package https://github.com/open2c/cooler. Saddle plots and compartment strength were calculated using the saddle plot functions in coolTools (https://cooltools.readthedocs.io/en/latest/notebooks/compartments_and_saddles.html). (Open2C et al., 2024). The bins with the strongest 20% of positive (for A) or negative (for B) PC1 values were used to calculate each interaction category (AA, BB).

#### Per-stage RNA-seq and Hi-C chromatin trajectory analysis

##### RNA-seq stage summary genes

The 69 canonical adipogenesis markers were organized into five sequential stages reflecting the temporal cascade of white adipocyte differentiation: Preadipocyte, Early, Mid/lipogenic, Late/terminal, and Lipolysis/metabolism. This grouping was chosen for two reasons.

First, the five stages align with distinct developmental transitions that can be independently interrogated: uncommitted precursor state (Preadipocyte); growth arrest and hormonal-cascade activation (Early); master adipogenic transcription factor induction and their direct lipogenic effector targets (Mid/lipogenic); mature adipocyte identity and lipid droplet formation (Late/terminal); and fully functional mature adipocyte metabolism (Lipolysis/metabolism). Second, this framework preserves the ability to detect stage-selective failures in the HGPS donors while remaining compact enough to visualize in a single figure panel. The genes used for each stage are described below along with their references.

ComBat-seq batch-corrected counts were normalized to counts per million (CPM). Replicates were averaged within each donor, time point, and condition, and each marker gene was z-score normalized across all time points. Stage scores in the graph are the mean of the z-scores of the markers in that stage; the progression index is the mean of late-(mean of preadipocyte + mean of early markers)/2.

##### Hi-C stage summary genes

To assess compartment-level chromatin dynamics at loci relevant to each stage of adipogenesis, we curated a panel of 71 canonical adipogenesis marker genes from published literature. We classified these genes into 5 categories: Preadipocyte, Early, Mid/lipogenic, Late/terminal, and Lipolysis and metabolism. Panel composition was defined by prior literature reports on each gene’s biological role in the adipogenic cascade rather than its expression pattern in the data set. For each stage, the mean compartment ΔPC value was calculated across all time points in each cell line. For day 0, mean eigenvector values across each stage were plotted. While the genes and references panel overlaps with the RNA-seq staging panel, additional references and their corresponding stage-specific genes are listed below:

#### For preadipocyte stage markers

DLK1/PREF1, PDGFRA, TCF21, CD34, CD44 [86–90], GATA2, GATA3, WNT10B [39], SFRP1 [91]

#### For early-stage markers

CEBPB, CEBPD, KLF15, KLF4, SREBF1, STAT5A, STAT5B, CREB1, EGR2 [92–100], FOXO1 [101], EBF1 [102], KLF5 [103]

#### For mid-stage/lipogenic markers

PPARG, CEBPA, RXRA, LPL, CD36, FASN, SCD, FABP4, GPD1, ACACA, DGAT1, DGAT2, ELOVL6, ME1, AGPAT2, LPIN1, INSR [104–120], PPARA, PPARD [121], ACACB [122], FABP5 [123], ACSL1 [124]

#### For late/terminal markers

LEP, PLIN1, PLIN4, PLIN5, CIDEC, ADIPOR1, ADIPOR2, RBP4, C1QTNF1, ZBTB16, TUSC5, NRIP1, FABP3, ADH1B [125–135], ADIPOQ [136], CIDEA [137], PLIN2 [138], AQP7 [139]

#### For lipolysis and metabolic markers

LIPE, PNPLA2, CFD, PCK1, PPARGC1A, LEPR, IRS1, IRS2, UCP2, UCP3, MGLL, ABHD5 [140–149], AKT2 [150], G0S2 [151]

### Genome-wide RNA-compartment concordance analysis

The per-cell-line gene tables (containing gene symbol and mean ΔPC across bins, as above) were joined by gene symbol to the D21 log2 fold-change from the RNA-seq master table for the same cell line. Only gene with both a non-zero mean ΔPC in that cell line and a threshold-passing D21 log2 fold-change entry were retained. Each retained gene was marked as concordant when the sign of the log2 fold-change and ΔPC were the same and discordant otherwise.

### Novel adipogenesis chromatin signature in the context of fibroblasts

Because fibroblasts remodel their chromatin landscape differently from MSCs, we looked for compartment features unique to fibroblast-associated adipogenesis. We identified 873 bins that switched from the B compartment in MSCs to the A compartment in fibroblasts and used them as a fibroblast-associated compartment signature. We further categorized and retained only bins that maintained A-compartment identity throughout induction in healthy fibroblast cell lines. Mapping these retained bins to GENCODE v19 gave us 515 protein-coding genes, of which 95 mapped to a curated adipogenesis panel (see Methods: Adipogenesis-focused concordance analysis) and 420 were unique to the fibroblast adipogenesis signature (Supplementary Figure 3D). From these 420 genes, we mapped the genes to the Enrichr GO terms.

## Data Availability

All RNA-seq and Hi-C sequencing data and processed data files are available at GEO accession number GSE351098. Adipogenesis differentiation images and quantification are available at Dryad: DOI: 10.5061/dryad.5x69p8dn5.

## Acknowledgements and Funding

We thank Dr. Amit Joshi, UTK, for advice about lipid droplet staining and analysis. We thank Tianchun Xue for help generating .mcool files. All members of the McCord lab provided helpful discussion and advice.

This work was funded by NIH NIGMS grant R35133557 to RPM and startup funds from Albert Einstein College of Medicine, Montefiore Einstein Comprehensive Cancer Center to RSM.

