## Supplementary Figures and Table for "Abnormal nuclear organization impairs adipogenic lineage commitment and terminal maturation in Hutchinson-Gilford progeria syndrome (HGPS)"

### Supplementary Figure 1

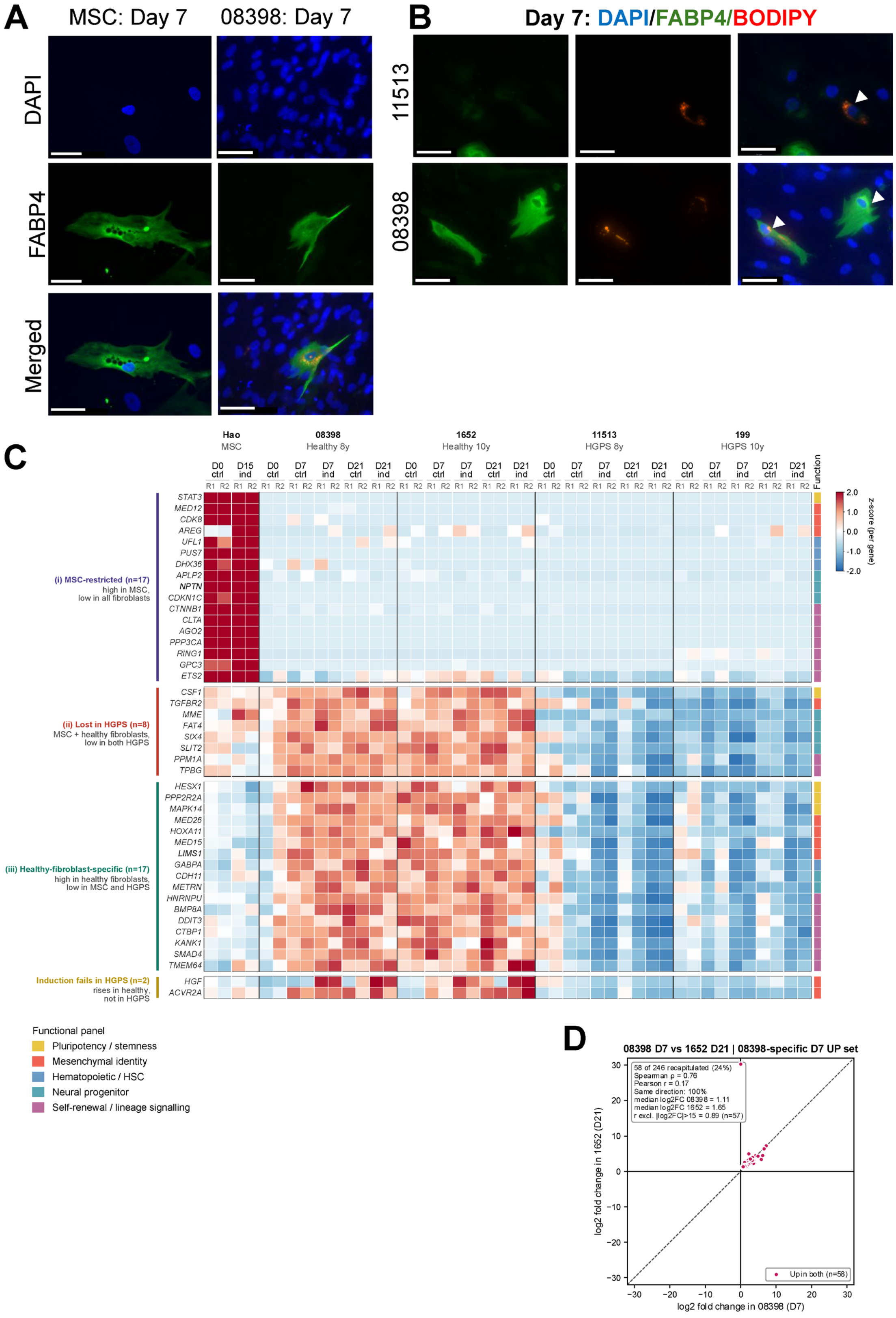

(A) Representative immunofluorescence images of bone marrow MSCs (MSC-PM) and healthy 8-year-old fibroblasts (08398) at day 7 (D7) of adipogenic induction, stained for FABP4 (green) and nuclei (DAPI, blue), Scale bars: 50 µm. (B) Representative images of progeria (11513) and healthy (08398) fibroblasts at D7, stained for nuclei (DAPI, blue), FABP4 (green), and neutral lipids (BODIPY, red). White arrowheads mark BODIPY+ lipid droplets. Lipid droplets can form without cage-like structures, so the two events can occur independently. Scale bars, 50 µm. (C) Heatmap of curated stem-cell markers in MSCs (Hao et al., D0 control and D15 induced) and in healthy (08398 and 1652) and progeria (11513 and 199) fibroblasts (D0 control, and D7 and D21 control and induced). Each column is one biological replicate (R1, R2). Expression values are z-scored per gene across all samples. Genes are grouped by expression patterns. (i) MSC-restricted (n=17), (ii) expressed in MSCs and healthy fibroblasts but lost in both HGPS lines (n=8), (iii) healthy fibroblast-specific (n=17), and (iv) induced in healthy fibroblasts but not in HGPS (n=2). The colored bar on the right shows each gene's functional category. (D) log2fold change of the 246 upregulated changes specifically in 08398 at D7, plotted against their log2fold change in 1652 at D21. Of these, 58 were also upregulated in 1652 at D21, all in the same direction. The dashed line marks y=x

### Supplementary Figure 2

**A**

| Timepoint | Number of shared genes in healthy | Retained in both HGPS donors | Retained in one HGPS donor | Lost in both HGPS donors | Direction discordant |
|---|---|---|---|---|---|
| D7 upregulated | 100 | 50% | 33% | 15% | 2% |
| D7 downregulated | 103 | 52% | 30% | 17% | 1% |
| D21 upregulated | 144 | 32% | 41% | 27% | NA |
| D21 downregulated | 163 | 34% | 43% | 22% | 1% |

**B**

Shared healthy: downregulated induced DEGs

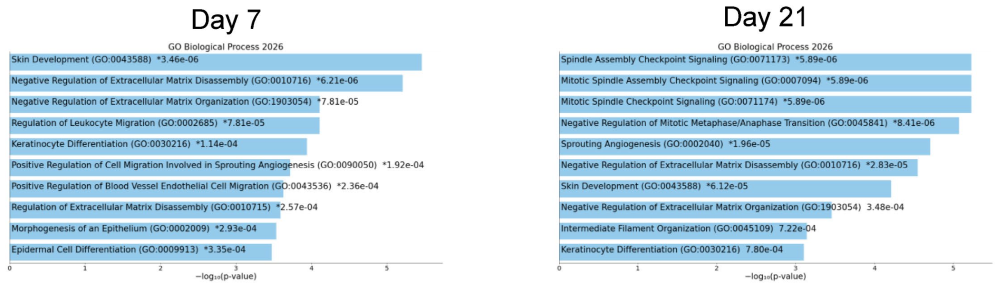


**C**

Shared progeria: downregulated induced DEGs

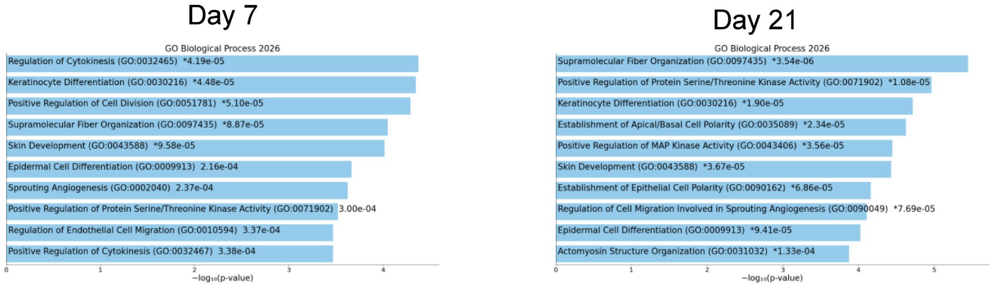


**D**

Shared healthy vs progeria: upregulated induced DEGs

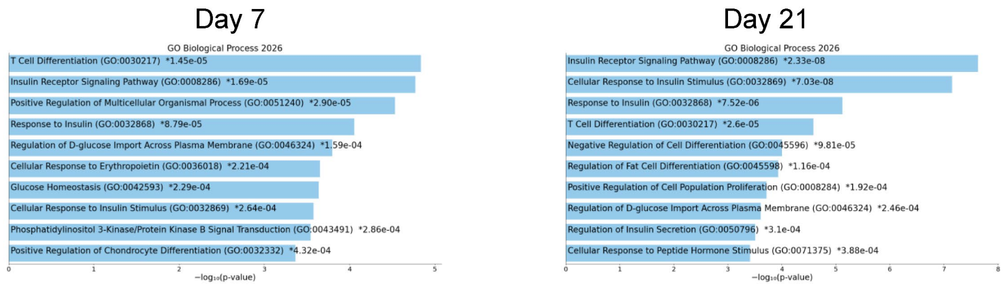


**E**

Shared healthy vs progeria: downregulated induced DEGs

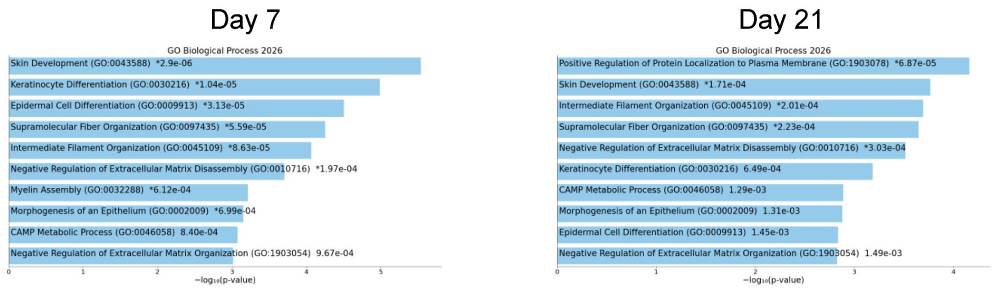

(A) Fate of DEGs shared by both healthy donors (08398 and 1652) and the two HGPS donors (11513 and 199), at D7 and D21, shown separately for upregulated and downregulated genes. For each set, the table gives the number of shared healthy DEGs and the percentages that were also differentially expressed in both HGPS donors (retained in both), in one HGPS donor only (retained in one), or neither (lost in both). The last column gives the percentage that changed in the opposite direction in HGPS. (B-E) GO biological process enrichment of shared DEG sets at D7 (left) and D21 (right): (B) downregulated DEGs shared by both healthy donors; (C) downregulated DEGs shared by both HGPS donors; (D) upregulated DEGs shared by healthy donors (08398 and 1652) in comparison to the progeria donors (11513 and 199); (E) downregulated DEGs shared by healthy donors (08398 and 1652) in comparison to the progeria donors (11513 and 199). The top 10 terms are ranked by -log10(p-value), and the p-value for each term is shown next to its bar. Enrichment analysis was performed using Enrichr (GO biological process 2026). DEGs were defined as adjusted $p<0.05$ (DESeq2) for adipogenic induction vs D0 controls.

### Supplementary Figure 3

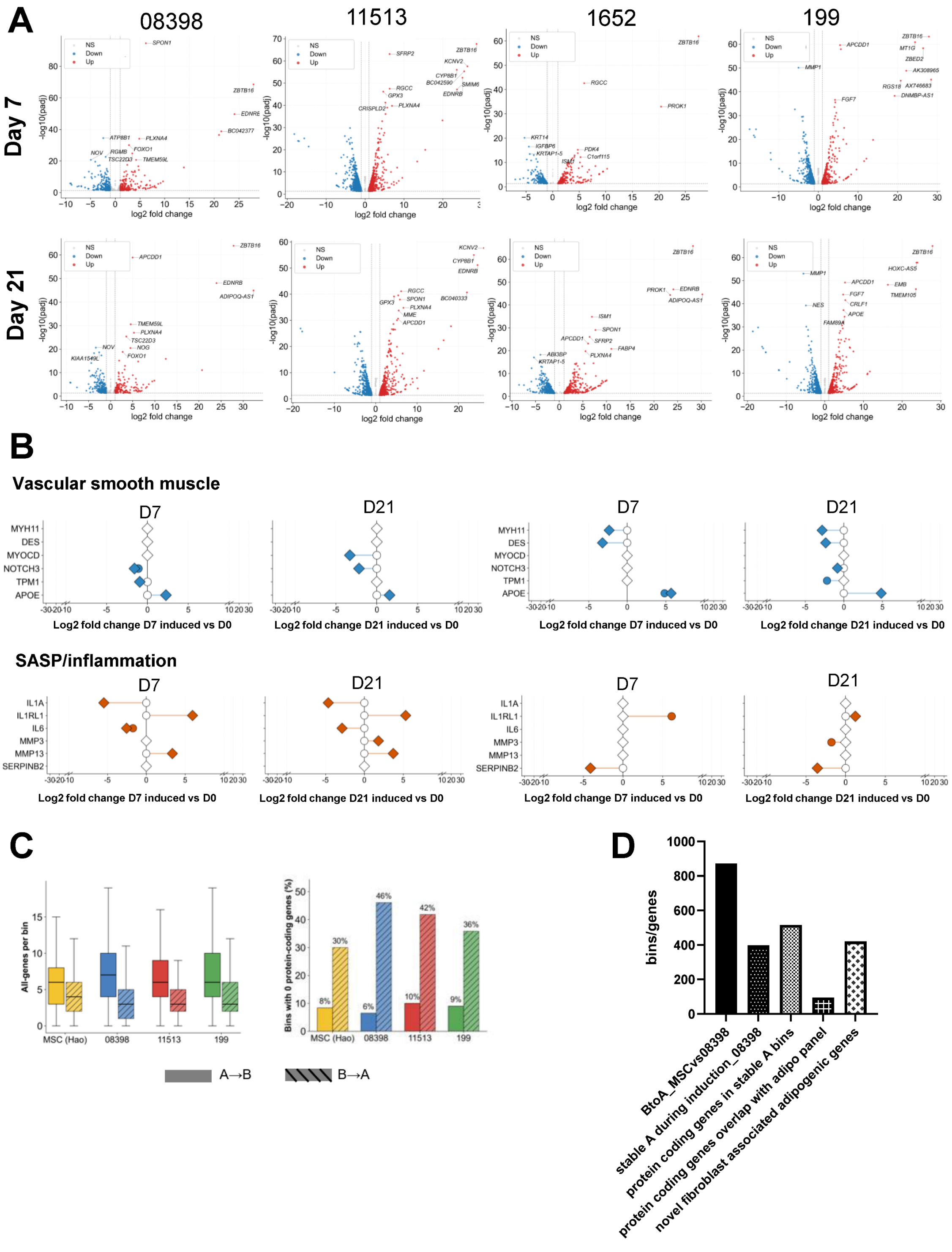

(A) Volcano plots of differential expression between adipogenic induction and D0 controls at D7 (top) and D21 (bottom), for healthy (08398, 1652) and progeria (11513, 199) fibroblasts. Age-matched pairs are shown side by side: 08398 with 11513 (8-year-old) and 1652 and 199 (10-year-old). Red marks upregulated genes and blue marks downregulated genes (adjusted p-value < 0.05, DESeq2); grey marks non-significant genes (NS). The most significant genes are labeled. Dashed lines show the significance thresholds (adjusted $p < 0.05$ and $|log2FC| \leq 1$). (B) Dumbbell plots of vascular smooth muscle genes (top) and SASP/inflammation genes (bottom) at D7 and D21, as in Figure 3A. The left plots compare 08398 (circles) with 11513 (diamonds), and the right plots compare 1652 with 199. Filled symbols mark DEGs; open symbols mark non-DEGs. (C) Gene content of compartment -switching bins in MSCs (Hao) and fibroblasts (08398, 11513, and 199), left: number of genes per 250-kb bin. Right: percentage of bins containing no protein-coding genes. Solid bars show A→B bins and hatched bars show B→A bins. Boxes show the median and interquartile range. (D) Derivation of the fibroblast-specific compartment signature. From left to right, the bars show: bins in the B compartment in MSCs but not in A (08398; n=873); the subset of these that stayed in the A compartment in 08398 (n= 395); the protein-coding genes in those stable A bins (n=515); the protein -coding genes that overlap the curated adipogenesis panel (n=95); and the remaining novel fibroblast-associated adipogenic genes (n=420).

#### Supplementary Figure 4

**A**

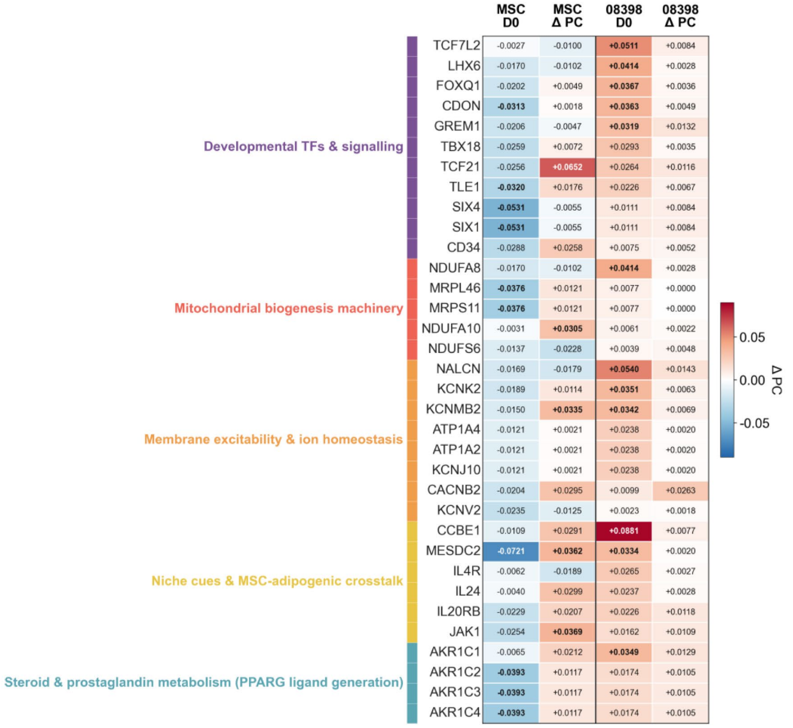


**B**

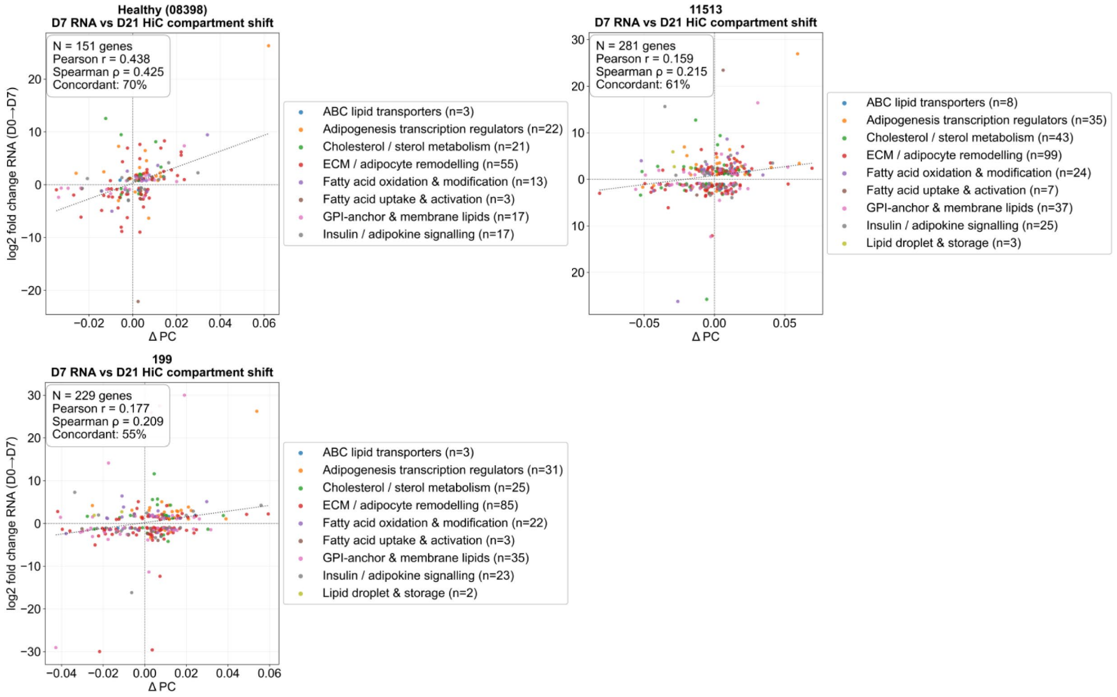

(A) Compartment eigenvector values for representative genes from the 420-gene fibroblast-specific adipogenic signature, grouped into five priming layers: developmental TFs and signaling; mitochondrial biogenesis machinery; membrane excitability and ion homeostasis; niche cues and MSC-adipogenic crosstalk; and steroid and prostaglandin metabolism. The columns show the D0 eigenvector and ΔPC in MSCs (D15) and in healthy (D21). Each gene takes the value of the 250-kb bin it falls in, so genes in the same bin share values. (B) RNA log2fold changes at D7 plotted against D21 compartment change for adipogenic-panel DEGs in 08398, 11513, and 199, colored by the functional category. The number of genes, Pearson and Spearman correlation coefficients, and the percentage concordance are shown in each plot. A gene is concordant when its expression and ΔPC change in the same direction. Dotted lines are shown as linear fits.

Supplementary Table 1: Hi-C sequencing and mapping statistics

| Sample | Total Raw Reads | Both Sides Mapped | %Both Sides Mapped | %Dangling Ends | Valid Pairs | Unique Valid Pairs | %Cis | %Cis >20 kb |
|---|---|---|---|---|---|---|---|---|
| GM08398_D0_R1 | 1046658487 | 30124988 | 2.88 | 31.11 | 13839846 | 6438139 | 88.11 | 29.53 |
| GM08398_D21 CONTROL_R1 | 655948121 | 239797839 | 36.56 | 9.09 | 205308278 | 146633946 | 81.18 | 55.67 |
| GM08398_D21 INDUCED ADIPOGENESIS_R1 | 654956585 | 235658531 | 35.98 | 7.42 | 208198369 | 149164252 | 81.04 | 59 |
| GM08398_D0_R2 | 580263643 | 202129194 | 34.83 | 5.85 | 183308162 | 134366826 | 79.56 | 56.02 |
| GM08398_D21 CONTROL_R2 | 599360465 | 269347888 | 44.94 | 13.2 | 198440802 | 113579145 | 84.43 | 47.68 |
| GM08398_D21 INDUCED ADIPOGENESIS_R2 | 442357138 | 188147781 | 42.53 | 12.11 | 144230593 | 92528862 | 83.81 | 51.98 |
| AG03199_D0_R1 | 627661893 | 237203888 | 37.79 | 9.37 | 194251905 | 125057640 | 80.94 | 53.25 |
| AG03199_D21 CONTROL_R1 | 1039151705 | 471361398 | 45.36 | 11.93 | 363506190 | 213293197 | 81.6 | 48.97 |
| AG03199_D21 INDUCED ADIPOGENESIS_R1 | 713504157 | 283727323 | 39.77 | 11.5 | 218458558 | 128908725 | 82.15 | 50.57 |
| AG03199_D0_R2 | 219791439 | 45343117 | 20.63 | 3.5 | 43052036 | 28489686 | 75.89 | 55.14 |
| AG03199_D21 CONTROL_R2 | 219791439 | 45343117 | 20.63 | 3.5 | 43052036 | 28489686 | 75.89 | 55.14 |
| AG03199_D21 INDUCED ADIPOGENESIS_R2 | 495220136 | 184099682 | 37.18 | 1.37 | 178763197 | 96625232 | 80.3 | 57.83 |
| AG11513_D0_R1 | 449682327 | 165572990 | 36.82 | 1.69 | 159811344 | 84794316 | 80.94 | 57.3 |
| AG11513_D21 CONTROL_R1 | 743791960 | 288449086 | 38.78 | 1.29 | 279911042 | 145660279 | 82.78 | 54.44 |
| AG11513_D21 INDUCED ADIPOGENESIS_R1 | 337703751 | 117549661 | 34.81 | 1.82 | 113674349 | 61073608 | 80.23 | 57.52 |
| AG11513_D0_R2 | 634439259 | 208416661 | 32.85 | 4.12 | 195198841 | 145744272 | 77.26 | 60.94 |
| AG11513_D21 CONTROL_R2 | 311554191 | 116882529 | 37.52 | 1.75 | 112604758 | 59590285 | 82.16 | 53.49 |
| AG11513_D21 INDUCED ADIPOGENESIS_R2 | 277153165 | 53955522 | 19.47 | 5.02 | 50012291 | 33034427 | 78.3 | 54.47 |
